# Anti-diuretic hormone ITP signals via a guanylate cyclase receptor to modulate systemic homeostasis in *Drosophila*

**DOI:** 10.1101/2024.02.07.579245

**Authors:** Jayati Gera, Marishia Agard, Hannah Nave, Austin B. Baldridge, Farwa Sajadi, Leena Thorat, Theresa H. McKim, Shu Kondo, Dick R. Nässel, Mitchell H. Omar, Jean-Paul V. Paluzzi, Meet Zandawala

## Abstract

Insects have evolved a variety of neurohormones that enable them to maintain their nutrient and osmotic homeostasis. While the identities and functions of various insect metabolic and diuretic hormones have been well-established, the characterization of an anti-diuretic signaling system that is conserved across most insects is still lacking. To address this, here we characterized the ion transport peptide (ITP) signaling system in *Drosophila*. The *Drosophila ITP* gene encodes five transcript variants which generate three different peptide isoforms: ITP amidated (ITPa) and two ITP-like (ITPL1 and ITPL2) isoforms. Using a combination of anatomical mapping and single-cell transcriptome analyses, we comprehensively characterized the expression of all three ITP isoforms in the nervous system and peripheral tissues. Our analyses reveal wide-spread expression of ITP isoforms. Moreover, we show that ITPa-producing neurons are activated and release ITPa during dehydration. Further, recombinant *Drosophila* ITPa inhibits diuretic peptide-induced renal tubule secretion *ex vivo*, thus confirming its role as an anti-diuretic hormone. Using a phylogenetic-driven approach, an *ex vivo* secretion assay and a heterologous mammalian cell-based assay, we identified and functionally characterized Gyc76C, a membrane guanylate cyclase, as a bona fide *Drosophila* ITPa receptor. Thus, recombinant ITPa application leads to increased cGMP production in HEK293T cells expressing *Drosophila* Gyc76C. Moreover, knockdown of Gyc76C in renal tubules abolishes the inhibitory effect of ITPa on diuretic hormone stimulated secretion. Extensive anatomical mapping of Gyc76C reveals that it is highly expressed in larval and adult tissues associated with osmoregulation (renal tubules and rectum) and metabolic homeostasis (fat body). Consistent with this expression, knockdown of Gyc76C in renal tubules impacts tolerance to osmotic and ionic stresses, whereas knockdown specifically in the fat body impacts feeding, nutrient homeostasis and associated behaviors. We also complement receptor knockdown experiments with *ITP* knockdown and ITPa overexpression in ITPa-producing neurons. Interestingly, the ITPa-Gyc76C pathway examined here is reminiscent of the atrial natriuretic peptide signaling in mammals. Lastly, we utilized connectomics and single-cell transcriptomics to identify synaptic and paracrine pathways upstream and downstream of ITPa-expressing neurons. Our analysis identifies pathways via which ITP neurons integrate hygrosensory inputs and interact with other homeostatic hormonal pathways. Taken together, our systematic characterization of ITP signaling establishes a tractable system to decipher how a small set of neurons integrates diverse inputs to orchestrate systemic homeostasis in *Drosophila*.

## Introduction

Metabolic and osmotic homeostasis are under strict control in organisms to ensure fitness and survival, as well as promote growth and reproduction. Homeostasis is achieved by regulatory mechanisms that impart plasticity to behaviors such as foraging, feeding, drinking, defecation, and physiological processes including digestion, energy storage/mobilization and diuresis. For any given homeostatic system, deviations from the optimal range are monitored by external and internal sensors. These in turn signal to central neuronal circuits where information about the sensory stimuli and internal states is integrated. The associated regulatory output pathways commonly utilize neuropeptides or peptide hormones to orchestrate appropriate behavioral and physiological processes (Rajan and Perrimon, 2011, Sternson, 2013, Jourjine *et al*., 2016, Lin *et al*., 2019, Nässel and Zandawala, 2019, Miroschnikow *et al*., 2020, Benevento *et al*., 2022, McKim *et al*., 2024). In mammals, hypothalamic peptidergic neuronal systems, in conjunction with peptide hormones released from the pituitary, are critical regulators of feeding, drinking, metabolic and osmotic homeostasis and reproduction (Sternson *et al*., 2013, Saper and Lowell, 2014, Le Tissier *et al*., 2017, Benevento *et al*., 2022). Several peptidergic pathways have also been delineated in insects that regulate similar homeostatic functions (Rajan and Perrimon, 2011, Schooley *et al*., 2012, Schoofs *et al*., 2017, Lin *et al*., 2019, Nässel and Zandawala, 2019, Nässel and Zandawala, 2020, Kim *et al*., 2021, Koyama *et al*., 2023). Some of these insect pathways originate in the neurosecretory centers of the brain and the ventral nerve cord (VNC), as well as in other endocrine cells located in the intestine (Raabe, 1989, Hartenstein, 2006, Zandawala *et al*., 2018a, Nässel and Zandawala, 2020, Zandawala *et al*., 2021, Koyama *et al*., 2023, McKim *et al*., 2024). Additionally, peptidergic interneurons distributed across the brain also play important roles in regulation of homeostatic behavior and physiology (Schlegel *et al*., 2016, Martelli *et al*., 2017, Lin *et al*., 2019, Yurgel *et al*., 2019, Miroschnikow *et al*., 2020, Nässel and Zandawala, 2020). Importantly, some insect neuropeptides are released by both interneurons and neurosecretory cells (NSC), indicating central and hormonal roles, respectively. One such example is the multifunctional ion transport peptide (ITP).

ITP derived its name from its first determined function in the locust *Schistocerca gregaria* where it increases chloride transport across the ileum and acts as an anti-diuretic hormone (Audsley *et al*., 1992b). Subsequent studies in other insects identified additional roles of ITP, including in reproduction, development, and post-ecdysis behaviors (Begum *et al*., 2009, Yu *et al*., 2016a). In *Drosophila,* ITP influences feeding, drinking, metabolism, and excretion (Galikova *et al*., 2018, Galikova and Klepsatel, 2022). Moreover, it has a localized interneuronal role in the *Drosophila* circadian clock system (Johard *et al*., 2009, Hermann-Luibl *et al*., 2014, Reinhard *et al*., 2024). The *Drosophila ITP* gene encodes five transcript variants, which generate three distinct peptide isoforms: one ITP amidated (ITPa) isoform and two ITP-like (ITPL1 and ITPL2) isoforms (Dircksen *et al*., 2008, Gramates *et al*., 2022) (Fig. 1A). ITPa is a C-terminally amidated 73 amino acid neuropeptide, while ITPL1 and ITPL2 are non-amidated and possess an alternate, extended C-terminus. ITPa and ITPL isoforms are also found in other insects, indicating conservation of the *ITP* splicing pattern (Dai *et al*., 2007). Moreover, insect ITP is homologous to crustacean hyperglycemic hormone (CHH) and molt inhibiting hormone (MIH), which together form a large family of multifunctional neuropeptides (Meredith *et al*., 1996, Dai *et al*., 2007, Drexler *et al*., 2007, Dircksen *et al*., 2008, Begum *et al*., 2009, Webster *et al*., 2012).

**Figure 1:**
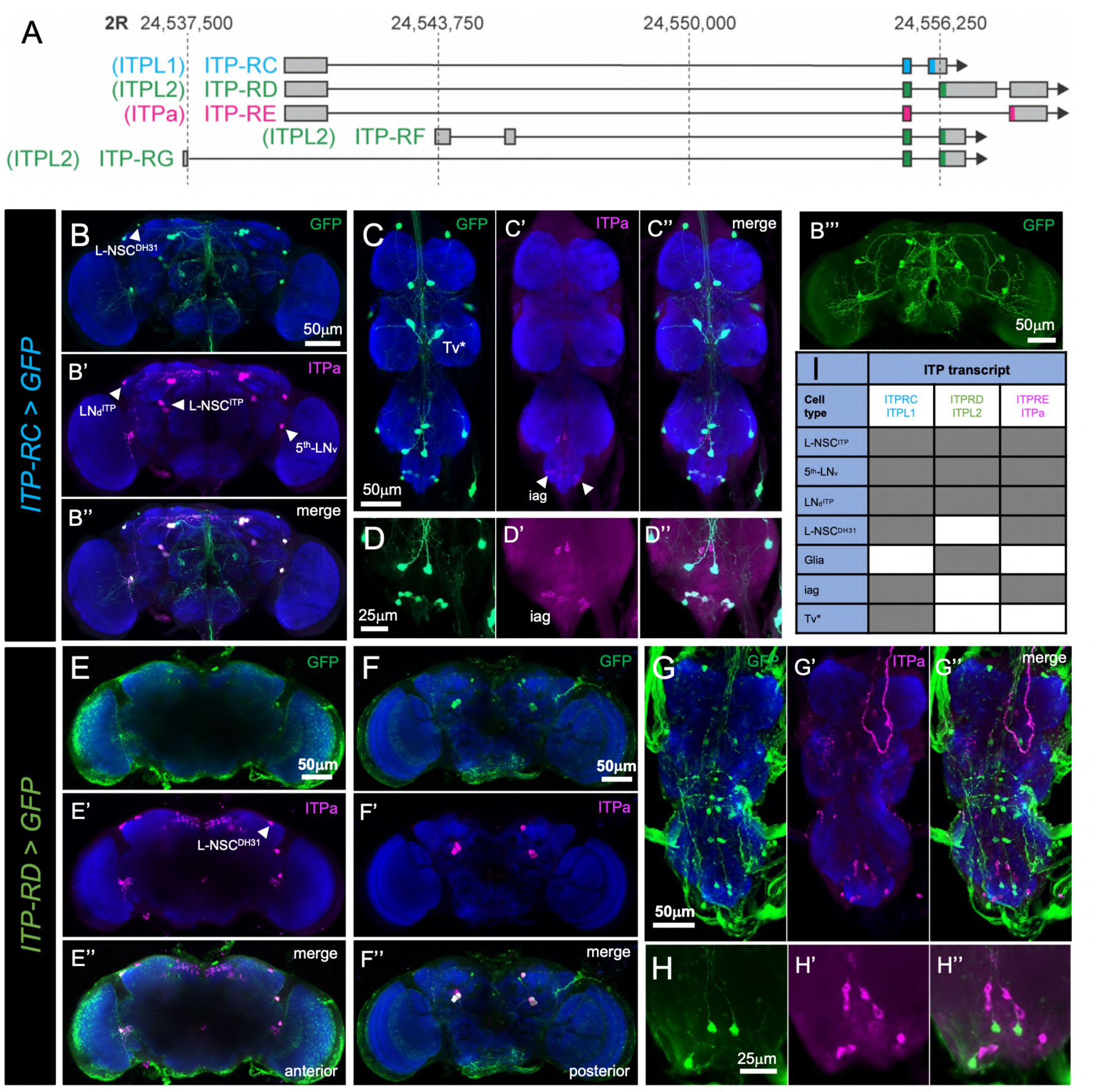
*ITP* splicing pattern and expression of *ITP* transcript variants in the nervous system of adult male *Drosophila*. **(A)** *Drosophila ITP* gene can generate 5 transcript variants (*ITP-RC*, *RD*, *RE*, *RF* and *RG*). *ITP-RC* encodes ITPL1 precursor, *ITP-RD*, *RF* and *RG* all encode ITPL2 precursor, and *ITP-RE* encodes a precursor that produces the amidated ITP (ITPa) peptide. Grey boxes represent exons and lines represent introns (drawn to scale). The regions encoding the open reading frame are colored (pink, green or blue). *ITP* is located on the second chromosome and numbers on the top indicate the genomic location. *ITP-RC-T2A-GAL4* drives GFP (UAS-JFRC81GFP) expression in the **(B)** brain and **(C and D)** ventral nerve cord (VNC). **B’’** shows another brain preparation (same as in Fig. 2A) where axons of ITP-RC neurons are clearly visible. All images are from male flies. Within the brain, ITP-RC is co-expressed with ITPa in four pairs of lateral neurosecretory cells (L-NSC^ITP^), one pair of diuretic hormone 31 (DH_31_)-expressing lateral neurosecretory cells (L-NSC^DH31^), one pair of 5^th^ ventrolateral neurons (5^th^-LN_v_) and one pair of dorsolateral neurons (LN_d_^ITP^). L-NSC^ITP^ and L-NSC^DH31^ are a subset of lateral neuroendocrine cells and the single pairs of 5^th^-LN_v_ and LN_d_^ITP^ belong to the circadian clock network. Within the VNC, ITP-RC is co-expressed with ITPa in abdominal ganglion neurons (iag), which innervate the rectal pad. In addition, ITP-RC is expressed in a pair of Tv* neurons near the midline in each thoracic neuromere. These neurons are located next to the FMRFamide-expressing Tv neurons (see Fig. 2B). *ITP-RD-T2A-GAL4* also drives GFP expression in the **(E and F)** brain and **(G and H)** VNC. ITP-RD is expressed in L-NSC^ITP^, 5^th^-LN_v_ and LN_d_^ITP^ neurons, as well as glia. Within the VNC, ITP-RD is expressed in neurons which are not iag or Tv* neurons. **(I)** Summary of ITP isoform expression within the nervous system. Grey box indicates presence and white box indicates absence.

ITPa is produced by a small set of interneurons and NSC in the *Drosophila* brain and VNC (Dircksen *et al*., 2008). While the expression of the ITPL peptides has not yet been investigated in *Drosophila*, studies in other insects indicate partial overlap with ITPa-expressing neurons (Drexler *et al*., 2007, Klocklerova *et al*., 2023). In order to delineate the targets of ITPa and ITPL and determine their modes of action, it is first necessary to identify, functionally characterize and localize the distribution of their cognate receptors. ITPa and ITPL receptors have been characterized in the silk moth *Bombyx mori* (Nagai *et al*., 2014). Surprisingly, *Bombyx* ITPa and ITPL were found to activate G-protein-coupled receptors (GPCRs) for pyrokinin and tachykinin neuropeptides, respectively (Nagai *et al*., 2014, Nagai-Okatani *et al*., 2016). Recently, ITPL2 was also shown to exert anti-diuretic effects via the tachykinin receptor 99D (TkR99D) in a *Drosophila* tumor model (Xu *et al*., 2023). Given the lack of structural similarity between ITPa/ITPL, pyrokinin and tachykinin, the mechanisms governing crosstalk between these diverse signaling pathways are still unclear. More importantly, the presence of any additional ITPa receptors in insects is so far unknown.

Here, we address these knowledge gaps by comprehensively characterizing ITP signaling in *Drosophila.* We used a combination of anatomical mapping and single-cell transcriptome analyses to localize expression of all three ITP isoforms in the nervous system and peripheral tissues. Importantly, we also functionally characterized the membrane-associated receptor guanylate cyclase, Gyc76C, and identified it as a *Drosophila* ITPa receptor. We show that ITPa-Gyc76C signaling to the fat body and renal tubules influences metabolic and osmotic homeostasis, respectively. Lastly, we identified synaptic and paracrine input and output pathways of ITP-expressing neurons using connectomics and single-cell transcriptomics, thus providing a framework to understand how ITP neurons integrate diverse inputs to orchestrate systemic homeostasis in *Drosophila*.

## Results

### Expression of ITP isoforms in the nervous system

In *Drosophila,* the *ITP* gene gives rise to five transcript variants: *ITP-RC*, *-RD*, -*RE*, *-RF* and *-RG*. *ITP-RC* encodes an ITPL1 precursor, *ITP-RD*, *-RF* and *-RG* all generate an ITPL2 precursor, whereas *ITP-RE* encodes a precursor which yields ITPa (Fig. 1A). Since the expression of *Drosophila* ITPL isoforms has not yet been mapped within the nervous system, we utilized specific T2A-GAL4 knock-in lines for ITPL1 (*ITP-RC-T2A-GAL4*) and ITPL2 (*ITP-RD-T2A-GAL4*) (Deng *et al*., 2019) to drive GFP expression. Concurrently, we stained these preparations using an antiserum against ITPa (Hermann-Luibl *et al*., 2014) to identify neurons co-expressing ITPa and ITPL isoforms (Fig. 1B-H).

In agreement with previous reports (Dircksen *et al*., 2008, Kahsai *et al*., 2010, Zandawala *et al*., 2018b), ITPa is localized in at least 7 bilateral pairs of neurons in the brain (Fig. 1B). Amongst these are 4 pairs of lateral NSC, L-NSC^ITP^ (also known as ipc-1 (Dircksen *et al*., 2008) or ALKs (de Haro *et al*., 2010)), that co-express ITPa, tachykinin, short neuropeptide F (sNPF) and leucokinin (LK) (Kahsai *et al*., 2010, Zandawala *et al*., 2018b). In addition, there is one pair each of dorsolateral neurons (LN_d_^ITP^) and 5^th^ ventrolateral neurons (5^th^-LN_v_), which are both part of the circadian clock network (Dircksen *et al*., 2008, Johard *et al*., 2009, Reinhard *et al*., 2024). Lastly, ITPa is weakly expressed in a pair of lateral NSC (also known as ipc-2a (Dircksen *et al*., 2008)). We demonstrate that these neurons (L-NSC^DH31^) co-express diuretic hormone 31 (DH_31_) (Fig. 1B and Fig. 2A). The L-NSC^DH31^ are also referred to as CA-LP neurons since they have axon terminations in the endocrine corpora allata (Siegmund and Korge, 2001, Kurogi *et al*., 2023). Interestingly, all ITPa brain neurons co-express ITPL1 (*ITP-RC*) (Fig. 1B). L-NSC^ITP^, possibly along with L-NSC^DH31^, are thus a likely source of ITPa and ITPL1 for hormonal release into the circulation.

**Figure 2:**
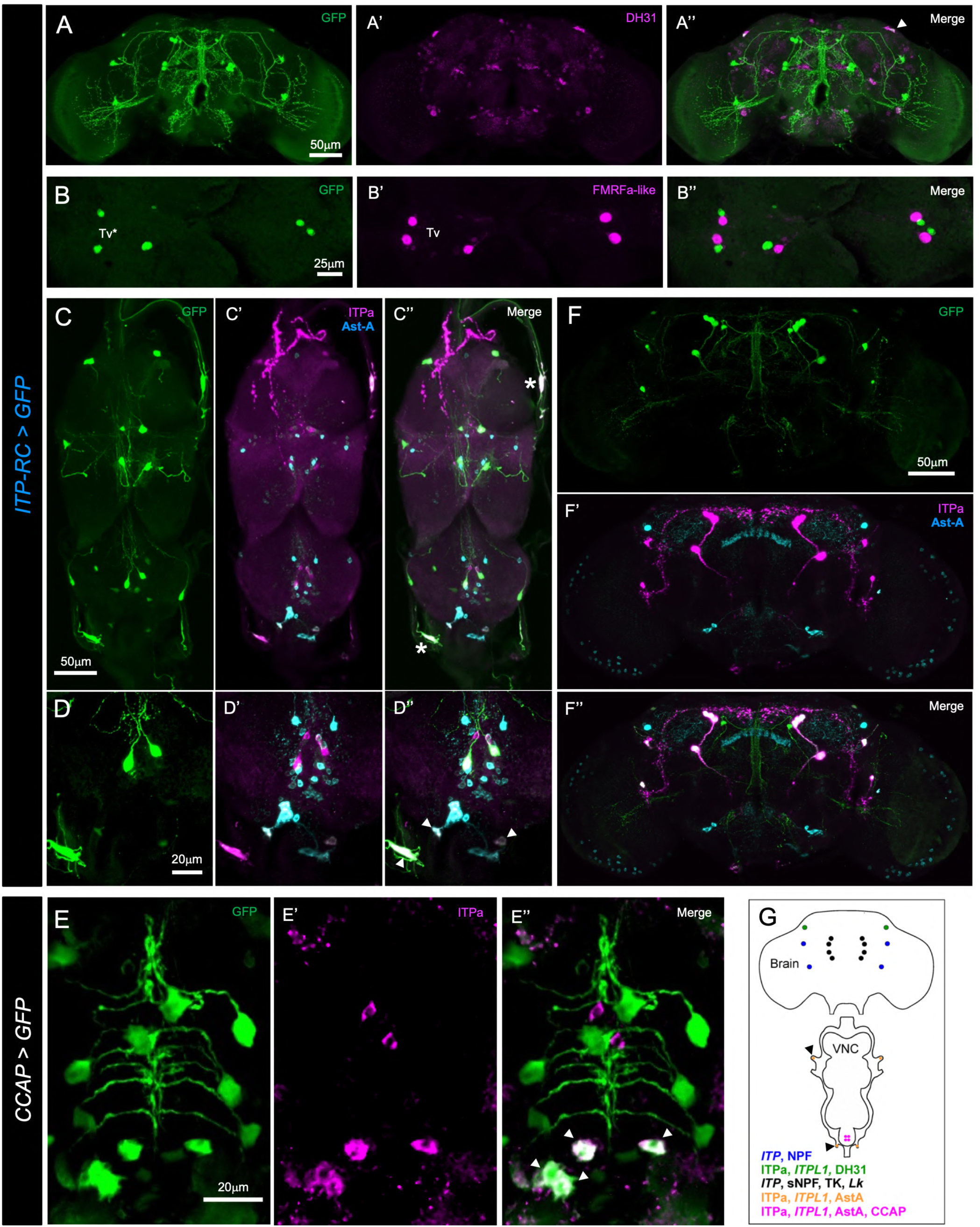
*ITP* is co-expressed with other neuropeptides in the nervous system of adult male *Drosophila*. **(A)** A single pair of *ITP-RC* > GFP-positive lateral neurosecretory cells in the dorsal brain (marked by an arrowhead) co-express diuretic hormone 31 (DH_31_). **(B)** *ITP-RC* drives GFP expression in a pair of Tv* neurons near the midline in each thoracic neuromere. These neurons are located next to the FMRFamide-expressing Tv neurons. **(C and D)** ITP-RC and ITPa-expressing peripheral neurons (marked by asterisk) and abdominal ganglion neurons (marked by arrowheads) co-express allatostatin-A (Ast-A) neuropeptide. **(E)** *CCAP* > GFP-positive neurons co-express ITPa (and Ast-A by extension) in the abdominal ganglion neurons (marked by arrowheads). **(F)** ITP-RC and ITPa-expressing neurons are distinct from Ast-A-expressing neurons in the brain. **(G)** Schematic of the nervous system showing neuropeptides (transcripts or mature peptides) expressed in ITP neurons. Peripheral neurons on one side are marked by arrowheads. Based on previous reports (Kahsai *et al*., 2010, Hermann-Luibl *et al*., 2014, Zandawala *et al*., 2018a) and the present study.

ITPa expression in the VNC is also sparse and is comprised of only the abdominal ganglion efferent neurons (iag) which innervate the hindgut and rectum (Fig. 1C, D) (Dircksen *et al*., 2008). In contrast, ITPL1 is expressed more widely, with *ITP-RC-T2A-GAL4* driven GFP detected in iag neurons as well as 14 additional neurons in the VNC (Fig. 1C, D). Six of these 14 neurons (Tv* in Fig. 1C) are located ventrally along the midline and closely resemble the six FMRFamide-expressing Tv neurons in the thoracic ganglia (Lundquist and Nässel, 1990, O’Brien *et al*., 1991). Our analysis reveals that the FMRFamide-expressing Tv neurons are distinct from the ITPL1-expressing ones, although their cell bodies are in close apposition (Fig.2B); hence, we refer to these ITPL1-expressing neurons as Tv*. Since abdominal ganglion efferent neurons that produce other neuropeptides have been described previously (Nässel and Zandawala, 2020), we asked whether ITPa/ITPL1-expressing iag neurons also express other neuropeptides. Interestingly, iag neurons co-express allatostatin-A (Ast-A) (Fig. 2C, D) and crustacean cardioactive peptide (CCAP) (Fig. 2E). In addition, peripheral neurons in the thoracic nerve roots also produce Ast-A and ITPa/ITPL1 (Fig. 2C, D); however, Ast-A and ITPa/ITPL1 are not co-expressed in the brain (Fig. 2F).

ITPL2 expression in the brain is also similar to ITPa and ITPL1 (Fig. 1E, F). However, *ITP-RD-T2A-GAL4* driven GFP was not detected in L-NSC^DH31^ but instead observed in glial cells surrounding the brain. In the VNC, ITPL2 was detected in peripheral glia as well as several neurons not producing ITPa and ITPL1 (Fig. 1G, H). Taken together, the three ITP isoforms exhibit partial overlapping distribution in the nervous system (Fig. 1I) and are, in some instances, also co-expressed with other neuropeptides in different subsets of neurons (Fig. 2G).

### Expression of ITP isoforms in peripheral tissues

In the silkworm *Bombyx mori* and the red flour beetle *Tribolium castaneum, ITP* gene products are also expressed outside the nervous system (Begum *et al*., 2009, Klocklerova *et al*., 2023). This peripheral source of ITP isoforms in *Bombyx* and *Tribolium* includes the gut enteroendocrine cells. In *Bombyx,* peripheral link neurons L1 which innervate the heart also express *ITP*. This prompted us to examine the expression of *Drosophila* ITP isoforms in tissues besides the nervous system. For this, we first examined the global expression of *ITP* using Fly Cell Atlas, a single-nucleus transcriptome atlas of the entire fly (Li *et al*., 2022). Surprisingly, this initial analysis revealed widespread expression of *ITP* across the fly (Fig. 3A-D). In particular, *ITP* is expressed in the trachea, Malpighian (renal) tubules (MTs), heart, fat body and gut (Fig. 3B-D). Fly Cell Atlas only provides expression levels for the entire gene, but not for individual transcript variants. Hence, we next mapped the cellular distribution of individual ITP isoforms in peripheral tissues using the T2A-GAL4 lines and ITPa-immunolabeling. As is the case in *Bombyx*, ITPL1 (*ITP-RC-T2A-GAL4* driven GFP) was detected in peripheral neurons that innervate the heart (Fig. 3E). In addition, axon terminals of iag neurons which innervate the rectum were also visible (Fig. 3F). No ITPL1 expression was observed in the fat body, midgut or MTs (Fig. 3G-I). Like ITPL1, ITPa immunoreactivity was also detected in a pair of peripheral neurons which innervate the heart and alary muscle (Fig. 3J), as well as in iag neuron axons that innervate the rectum (Fig. 3K). ITPa immunoreactivity was not detected in the midgut (Fig. 3L). In comparison to ITPa and ITPL1, ITPL2 is more broadly expressed in peripheral tissues. Thus, *ITP-RD-T2A-GAL4* drives GFP expression in the heart muscles and the neighboring pericardial nephrocytes (Fig. 3M), as well as in cells of the middle midgut (Fig. 3N), posterior midgut (Fig. 3O), ureter (Fig. 3P) and trachea (Fig. 3Q), but not the fat body (Fig. 3R). In summary, Fly Cell Atlas data are largely in agreement with our comprehensive anatomical mapping of individual ITP isoforms. The widespread expression of *ITP* in peripheral tissues can be largely attributed to ITPL2. Expression of ITPL1 and ITPa, on the other hand, is more restricted and overlaps in cells innervating the heart and rectum.

**Figure 3:**
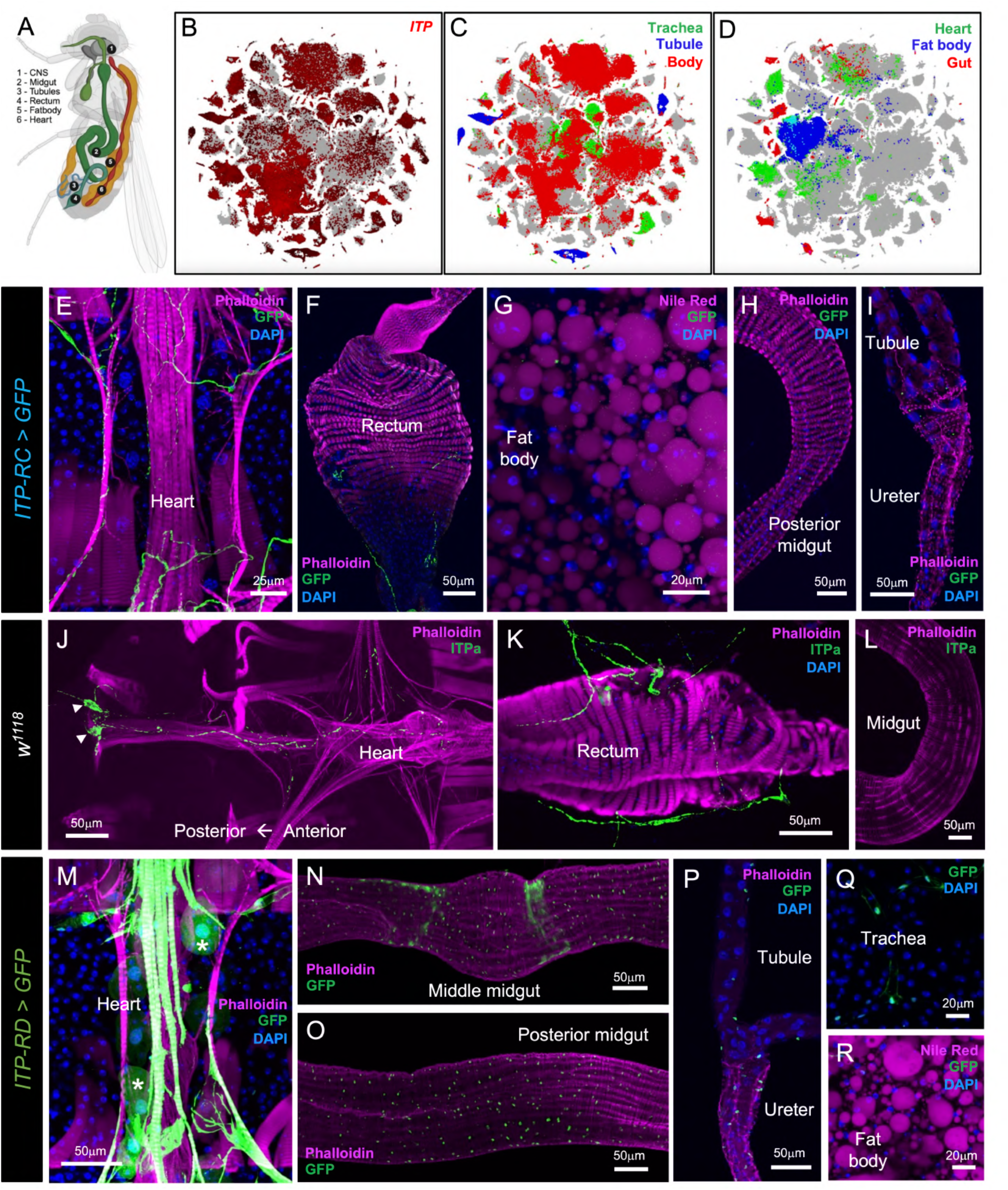
*ITP* expression in peripheral tissues of adult male *Drosophila.* **(A)** Schematic showing the location of tissues where *ITP* is expressed. Created with BioRender.com. **(B)** t-SNE visualization of single-cell transcriptomes showing *ITP* expression in different tissues of adult *Drosophila. ITP* is broadly expressed in peripheral tissues, including **(C)** trachea, Malpighian tubules (tubule), body, **(D)** heart, fat body and gut. *ITP-RC-T2A-GAL4* drives GFP (UAS-JFRC81GFP) expression in **(E)** peripheral neurons with axons innervating the heart and **(F)** abdominal ganglion neurons which innervate the rectum. ITP-RC is not expressed in **(G)** the fat body, **(H)** midgut or **(I)** Malpighian tubules. ITPa immunolabelling is present in **(J)** a pair of peripheral neurons (cell bodies marked by arrowheads) innervating the heart and **(K)** in abdominal ganglion neurons which innervate the rectum, but **(L)** absent in the midgut. *ITP-RD-T2A-GAL4* drives GFP expression in **(M)** the heart and nephrocytes (marked by asterisk), **(N)** middle midgut, **(O)** posterior midgut, **(P)** ureter and **(Q)** trachea. **(R)** ITP-RD is not expressed in the fat body.

### Identification of Gyc76C as a putative ITP receptor

ITP has been shown to influence osmotic, ionic and metabolic homeostasis in insects, including *Drosophila* (Audsley *et al*., 1992b, Galikova *et al*., 2018, Galikova and Klepsatel, 2022). Considering that the control of hydromineral balance requires stringent integration of all excretory organs, including the rectum and MTs in adult flies, we hypothesized that a putative *Drosophila* ITP receptor would be expressed in these tissues. Our expression mapping of ITP isoforms suggests that osmotic/ionic homeostasis is regulated, at least in part, via a direct effect on the rectum, which is responsible for water and ion reabsorption (Phillips *et al*., 1987, Coast *et al*., 2002, O’Donnell, 2008). Additionally, we also expect a putative ITP receptor to be expressed in the fat body, which is a major metabolic tissue. First, we explored if the *Drosophila* orthologs of *Bombyx* ITPa and ITPL receptors (Fig. 4 Supplement 1A) could also function as ITPa/ITPL receptors in *Drosophila* by examining their expression in the gut, fat body and MTs. *Bombyx* ITPa activates two GPCRs, which are orthologous to *Drosophila* pyrokinin 2 receptor 1 (PK2-R1) and an orphan receptor (CG30340) whose endogenous ligand in *Drosophila* is still unknown (Nagai *et al*., 2014). In addition, *Bombyx* ITPL and tachykinin both activate another GPCR which is related to the *Drosophila* tachykinin receptor at 99D (TkR99D) (Fig. 4 Supplement 1A) (Nagai *et al*., 2014, Nagai-Okatani *et al*., 2016). Our analysis revealed that neither of the three candidate *Drosophila* receptors, PK2-R1, TkR99D and CG30340, are expressed in the epithelial cells of the rectal pad which mediate ion and water reabsorption (Fig. 4 Supplement 1B-D). GFP expression for PK2-R1 and TkR99D was observed in axons innervating the rectum, suggesting that they are expressed in efferent neurons in the abdominal ganglion. In addition, all three receptors were expressed in the midgut or in neurons innervating it (Fig. 4 Supplement 1E-G), indicating that the GAL4 drivers and the GFP constructs used here are strong enough to report expression in other tissues. Further, single-nucleus RNA sequencing analyses revealed that neither PK2-R1 nor CG30340 are expressed in the cells of the fat body (Fig. 4 Supplement 1H) and MTs (Fig. 4 Supplement 1I). TkR99D, on the other hand, is not detected in the fat body (Fig. 4 Supplement 1H) but is expressed in the MT stellate cells (Fig. 4 Supplement 1I), where it mediates diuretic actions of tachykinin (Agard *et al*., 2024). Hence, expression mapping and/or previous functional analysis of PK2-R1, CG30340 and TkR99D indicates that they are not suited to mediate the anti-diuretic and metabolic effects of ITPa/ITPL. To test this experimentally, we generated recombinant ITPa for analysis of GPCR activation *ex vivo*. We found that recombinant ITPa failed to activate both TkR99D (Fig. 4 Supplement 1J) and PK2-R1 (Fig. 4 Supplement 1K) heterologously expressed in mammalian CHO-K1 cells. As a control, we showed that their natural respective ligands, tachykinin 1 and pyrokinin 2, resulted in strong receptor activation (Fig. 4 Supplement 1J, K). Taken together, these experiments suggest that receptor(s) for *Drosophila* ITPa appear to be evolutionary divergent from *Bombyx* ITPa/ITPL receptors.

**Figure 4:**
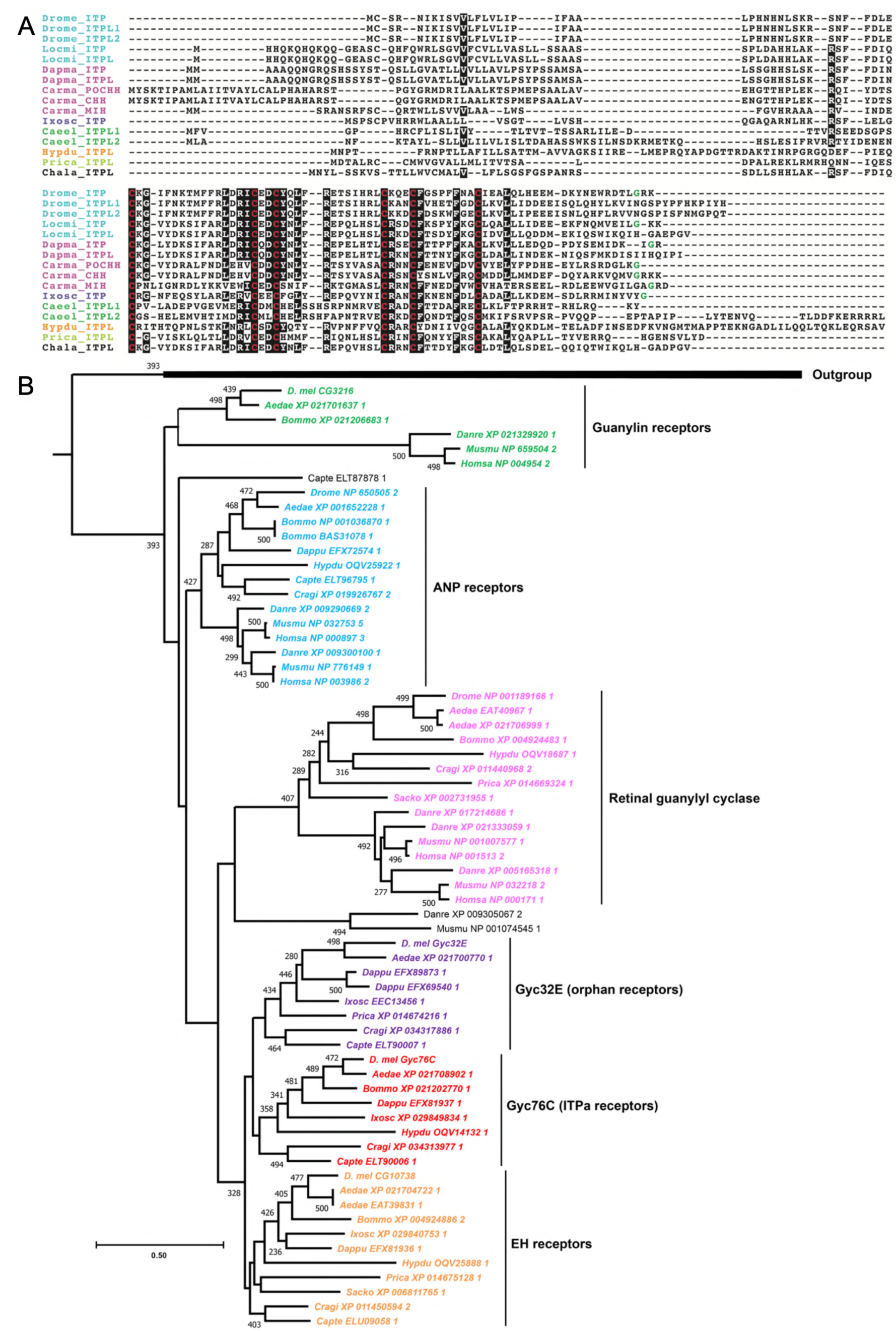
ITP signaling components are found in protostomes. **(A)** Multiple sequence alignment of ITP precursor sequences. ITP is homologous to crustacean hyperglycemic hormone (CHH) and molt-inhibiting hormone (MIH). Note the conservation of six cysteine residues (highlighted in red) across all the species. C-terminal glycine which is predicted to undergo amidation is colored in green. Species abbreviations: Drome, *Drosophila melanogaster*; Locmi, *Locusta migratoria*; Dapma, *Daphnia magna*; Carma, *Carcinus maenas*; Ixosc, *Ixodes scapularis*; Caeel, *Caenorhabditis elegans*; Hypdu, *Hypsibius dujardini*; Prica, *Priapulus caudatus*; Chala, *Charonia lampas*. **(B)** Maximum-likelihood phylogeny of membrane guanylate cyclase receptors identifies two clades that are restricted to protostome phyla which also have ITP. The clade containing *D. melanogaster* Gyc76C receptor are the putative ITPa receptors. Bootstrap values higher than 200 (based on 500 replicates) are indicated adjacent to the nodes. *Drosophila* guanylate cyclase alpha and beta subunits were used as outgroups.

Having ruled out the *Bombyx* ITPa/ITPL receptor orthologs as potential candidates, we next employed a phylogenetic-driven approach to identify additional novel ITP receptor(s) in *Drosophila* and other species. Since neuropeptides and their cognate receptors commonly coevolve (Park *et al*., 2002, Jekely, 2013), we reasoned that the phyletic distribution of ITP would closely mirror that of a putative ITP receptor. Hence, we first used BLAST and Hidden Markov Model (HMM)-based searches to identify *ITP* genes across all animals. Our analyses retrieved *ITP/CHH/MIH*-like genes in arthropods, nematodes, tardigrades, priapulid worms and mollusks (Fig. 4A). A comparison of representative ITP precursor sequences from different phyla reveals that the six cysteines and a few amino acid residues adjacent to them are highly conserved (Fig. 4A). Thus, *ITP* appears to be restricted to protostomian invertebrates and does not have orthologs in deuterostomian invertebrates and vertebrates. To identify putative orphan receptor(s) which follow a similar phyletic distribution, we performed a phylogenetic analysis of receptors from different vertebrate and invertebrate phyla. We specifically focused on membrane guanylate cyclase receptors (mGC) that all couple with the cGMP pathway because ITP/CHH stimulation has previously been shown to result in an increase in cGMP (Dircksen, 2009, Nagai *et al*., 2014). Phylogenetic analysis grouped mGC into six distinct clades (Fig. 4B). Four of these comprise guanylin, atrial natriuretic peptide (ANP), retinal guanylyl cyclase and eclosion hormone receptors. Importantly, we retrieved two clades which only contain receptors from protostomian invertebrates (Fig. 4B). One clade includes *Drosophila* Gyc76C and another includes Gyc32E. Both receptors meet the peptide-receptor co-evolution criteria for ITP receptor identification. However, single-nucleus sequencing data indicate that *Gyc76C* is more highly expressed than *Gyc32E* in MTs (Fig. 4 Supplement 2A). Independently, we did not detect *Gyc32E-GAL4* driven GFP expression in MTs (Fig. 4 Supplement 2B) and rectal pad (Fig. 4 Supplement 2C) but it was present in the hindgut (Fig. 4 Supplement 2C) and fat body (Fig. 4 Supplement 2D). *Gyc32E-GAL4* is also expressed in a subset of insulin-producing cells (IPCs; labelled with antibody against DILP2) in the brain (Fig. 4 Supplement 2E). Thus, the lack of *Gyc32E* expression in osmoregulatory tissues, coupled with the fact that Gyc76C was previously implicated in the ITP signaling pathway in *Bombyx* (Nagai *et al*., 2014), prompted us to focus on Gyc76C further.

### Gyc76C expression in Drosophila

If Gyc76C functions as an ITP receptor in *Drosophila,* it should be expressed in cells and tissues that are innervated by ITPa/ITPL-expressing neurons as well as in tissues which mediate some of the known hormonal functions of ITP. To validate this prediction, we used a recently generated T2A-GAL4 knock-in line for Gyc76C (Fig. 5A) (Kondo *et al*., 2020) and used it to comprehensively map Gyc76C expression throughout larval *Drosophila* (Fig. 5 Supplement 1) and in adult males (Fig. 5B-O) and females (Fig. 5 Supplement 2). In males, *Gyc76C-T2A-GAL4* drives GFP expression throughout the adult intestinal tract, including the anterior midgut (Fig. 5B), ureter (of renal tubules) (Fig. 5C) and posterior midgut (Fig. 5D). Importantly, in agreement with the role of *Drosophila* ITP in regulating osmotic (Galikova *et al*., 2018) and metabolic homeostasis (Galikova and Klepsatel, 2022), Gyc76C is highly expressed in the renal tubules (Fig. 5E), rectum (Fig. 5F) and adipocytes of the fat body (Fig. 5G). Moreover, we see a convergence of ITPa-immunolabeled axon terminations and Gyc76C expression in the anterior midgut (Fig. 5H) and the rectal papillae in the rectum (Fig. 5I), the latter of which are important for water reabsorption as first proposed nearly a century ago (Wigglesworth, 1932). Gyc76C is also broadly expressed in neurons throughout the brain (Fig. 5J) and VNC (Fig. 5K). Consistent with the role of ITP in regulating circadian rhythms, Gyc76C is expressed in glia clock cells (Fig. 5L) and subsets of dorsal clock neurons (labelled with antibody against the clock protein Period) that are near the axon terminations of the clock neurons LNd^ITP^ and 5^th^-LN_v_ (Fig. 5M). Gyc76C is not expressed in lateral clock neurons which are situated more closely to ITPa-expressing clock neurons (Fig. 5L). Similar to males, *Gyc76C-T2A-GAL4* also drives GFP expression in the female fat body (Fig. 5 Supplement 2A), renal tubules (Fig. 5 Supplement 2B), midgut (Fig. 5 Supplement 2C), brain (Fig. 5 Supplement 2D), VNC (Fig. 5 Supplement 2E) and subsets of dorsal clock neurons (Fig. 5 Supplement 2F, G). Interestingly, Gyc76C is not expressed in male IPCs (labelled with antibody against DILP2) (Fig. 5N) but is expressed in a subset of female IPCs (Fig. 5 Supplement 2H). However, Gyc76C is not expressed in endocrine cells producing glucagon-like adipokinetic hormone (AKH) in either males (Fig. 5O) or females (Fig. 5 Supplement 2I). Interestingly, L-NSC^DH31^, which innervate the corpora allata, might utilize both DH_31_ and ITPa/ITPL1 to modulate juvenile hormone production since Gyc76C is expressed in the corpora allata (Fig. 5O). Lastly, we also explored the distribution of Gyc76C in larval tissues (Fig. 5 Supplement 1), where expression was detected in the adipocytes of the fat body, all the regions of the gut and in renal tubules (Fig. 5 Supplement 1A-H). Gyc76C is widely distributed in the larval nervous system, with high expression in the endocrine ring gland, where ITPa-immunoreactive axons terminate (Fig. 5 Supplement 1I). Taken together, the cellular expression of Gyc76C in the nervous system and peripheral tissues of both larval and adult *Drosophila* further indicates that it could mediate the known effects of ITP.

**Figure 5:**
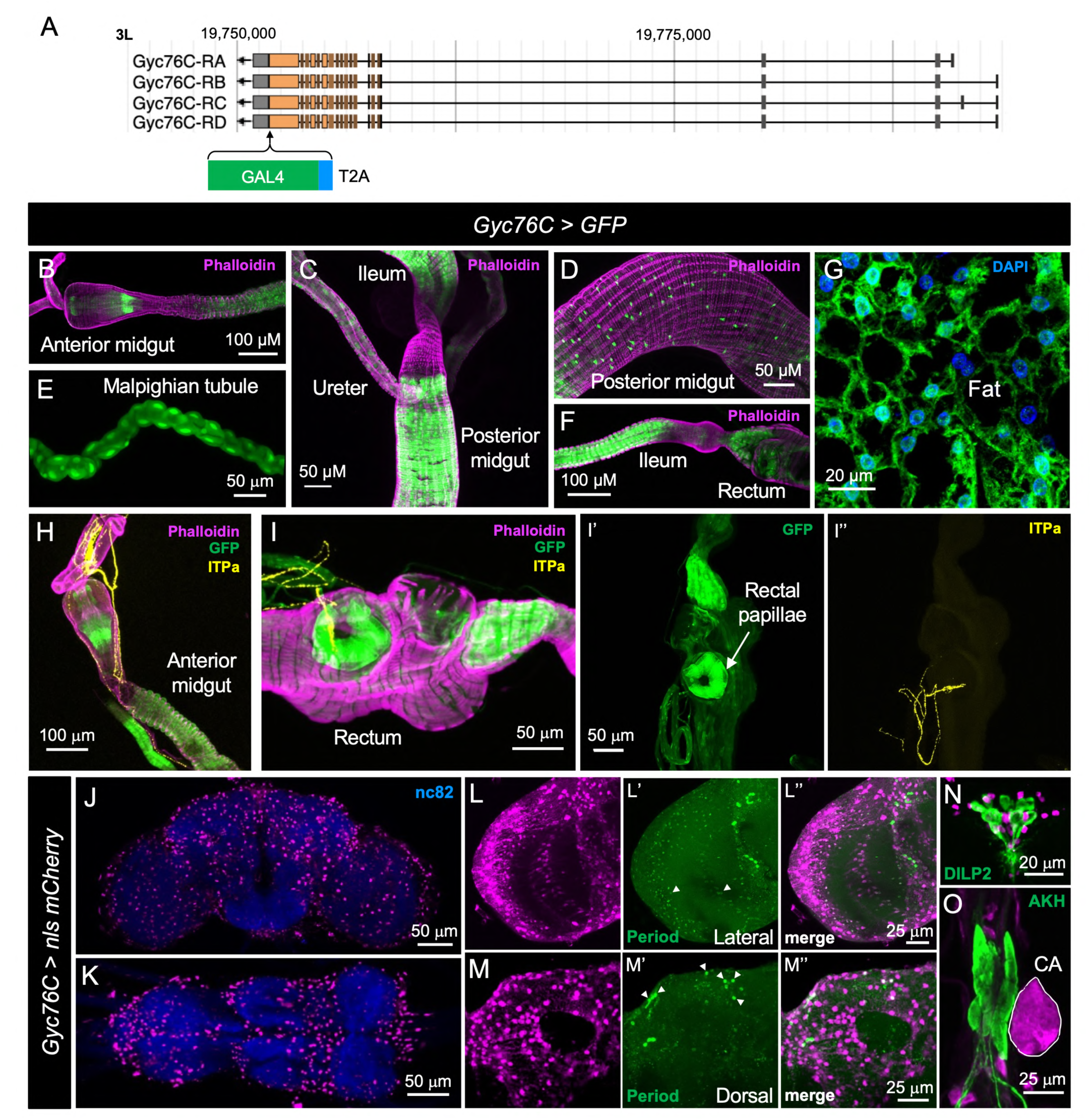
Gyc76c expression in adult male *Drosophila*. **(A)** Schematic showing the generation of *Gyc76C-T2A-GAL4* knock-in line. *Gyc76C-T2A-GAL4* drives GFP (UAS-JFRC81GFP) expression in the **(B)** anterior midgut, **(C)** ureter, **(D)** posterior midgut, **(E)** Malpighian tubules, **(F)** ileum, rectum, **(G)** and adipocytes in the fat body. Gyc76C is expressed in the regions of **(H)** the anterior midgut and **(I)** rectal papillae in the rectum that are innervated by ITPa-expressing neurons. Gyc76C is also broadly expressed in the **(J)** brain and **(K)** ventral nerve cord. **(L)** Gyc76C is expressed in glial clock cells and **(M)** subsets of dorsal clock neurons (both labelled by Period antibody and marked by arrowheads). Gyc76C is not expressed in **(N)** insulin-producing cells (labelled by DILP2 antibody) and **(O)** adipokinetic hormone (AKH) producing endocrine cells but is expressed in the corpora allata (CA) (marked in white).

### Gyc76C is necessary for ITPa-mediated inhibition of renal tubule secretion ex vivo

ITP has been shown to modulate osmotic homeostasis in *Drosophila* by suppressing excretion (Galikova *et al*., 2018). While the precise mechanisms underlying the anti-diuretic effects of *Drosophila* ITP are not known, previous research in other systems provide important insights. For instance, in the locust *Schistocerca gregaria*, ITPa but not ITPL promotes ion and water reabsorption across the hindgut (Audsley *et al*., 1992a, Audsley *et al*., 1992b, King *et al*., 1999, Wang *et al*., 2000), thereby promoting anti-diuresis. Previous experiments have shown that the MTs are also targeted by anti-diuretic hormones: CAPA neuropeptides inhibit diuresis in some insects including *Drosophila* via direct hormonal actions on the renal tubules (Paluzzi *et al*., 2008, MacMillan *et al*., 2018, Sajadi *et al*., 2020, Sajadi *et al*., 2023). Given the expression of Gyc76C, our candidate ITP receptor, in both the hindgut and renal tubules, ITP could modulate osmotic and/or ionic homeostasis by targeting these two excretory organs. Hence, we utilized the Ramsay assay (Fig. 6A) to monitor *ex vivo* fluid secretion by MTs in response to application of recombinant ITPa. Interestingly, recombinant ITPa does not influence rates of secretion by unstimulated tubules (Fig. 6B). Since the basal secretion rates are quite low, we tested if ITPa can inhibit secretion stimulated by LK, a diuretic hormone targeting stellate cells (O’Donnell *et al*., 1996), and a calcitonin-related peptide, DH_31_, which acts on principal cells (Johnson *et al*., 2005). Recombinant ITPa inhibits LK-stimulated secretion by MTs from *w^1118^* flies (Fig. 6C), indicating that stellate cell driven diuresis is sensitive to this anti-diuretic hormone. Similarly, DH_31_-stimulated secretion was also inhibited by recombinant ITPa (Fig. 6D), demonstrating that principal cells are also modulated by ITPa. This result confirmed that the effect of ITPa on osmotic homeostasis are mediated, at least partially, via actions on renal tubules and that both major cell types are targeted.

**Figure 6:**
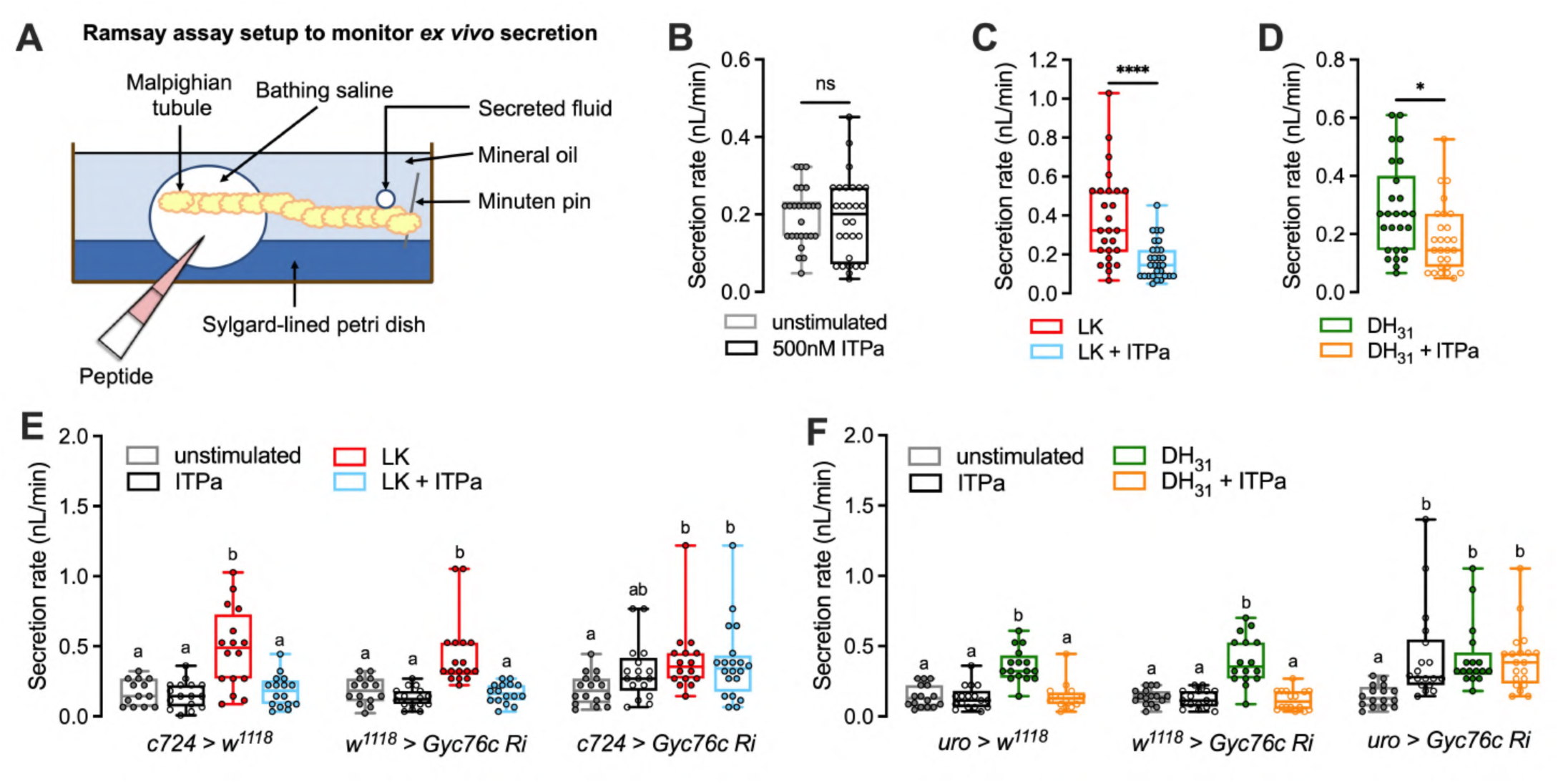
Recombinant *Drosophila* ITPa inhibits Malpighian tubule secretion via Gyc76C. **(A)** Schematic of Ramsay assay used to monitor *ex vivo* secretion by tubules. **(B)** Application of *Drosophila* 500nM ITPa does not affect basal secretion rates by unstimulated tubules. 500nM ITPa inhibits both **(C)** 10nM leucokinin (LK)-stimulated and **(D)** 1μM diuretic hormone 31 (DH_31_)-stimulated secretion rates. Importantly, while 500nM ITPa inhibits **(E)** 10nM LK-stimulated secretion and **(F)** 1μM DH31-stimulated by renal tubules from control flies, this inhibitory effect is abolished in tubules where *Gyc76C* has been knocked down with *UAS-Gyc76C RNAi* (*#106525*) in stellate cells using the *c724-GAL4* and in principal cells using *uro-GAL4*. Male Malpighian tubules were used for all experiments. For **B-D,**\* p < 0.05 and **** p < 0.0001 as assessed by unpaired *t* test. For **E and F**, within each genotype, different letters denote secretion rates that are significantly different from one another (p < 0.05) as assessed by two-way ANOVA followed by Tukey’s multiple comparisons test.

We next utilized the Ramsay assay to assess if Gyc76C is necessary for the inhibitory effects of ITPa on renal tubule secretion. Notably, knocking down expression of Gyc76C in stellate cells using *c724-GAL4* abolished the anti-diuretic action of ITPa in LK-stimulated tubules (Fig. 6E). Similarly, tubules in which *Gyc76C* was knocked down using the *LK receptor GAL4* (Zandawala *et al*., 2018b) do not exhibit reduced secretion following ITPa application (Fig. 6 Supplement 1). Additionally, knocking down expression of Gyc76C in principal cells using *uro-GAL4* abolished the anti-diuretic action of ITPa in DH_31_-stimulated tubules (Fig. 6F). Surprisingly, application of ITPa alone promotes fluid secretion compared to unstimulated controls in tubules with Gyc76C knockdown (Fig. 6E, F). This effect is more prominent in tubules with Gyc76C knockdown specifically in the principal cells (Fig. 6E, F). This suggests that ITPa could also interact with other yet unknown receptors in addition to Gyc76C. Nonetheless, these results indicate that ITPa exerts its anti-diuretic effects on MTs via Gyc76C, which acts as a functional ITPa receptor in both stellate and principal cells of the tubules.

### ITPa activates Gyc76C in HEK293T cells

To further test whether ITPa activates Gyc76C, we leveraged a heterologous expression assay. HEK293T cells were transfected with Gyc76C-HA and/or the fluorescent protein-based cGMP indicator Green cGull (Matsuda *et al*., 2017) and subjected to live-cell imaging (Fig. 7A). At 4 minutes after 50nM, 250nM, or 500nM ITPa treatment, cells expressing HA-tagged Gyc76C and Green cGull exhibited 34%, 88.7%, and 111.4% increases in fluorescence intensity, respectively, indicating a dose-dependent increase in cGMP (Fig. 7B-E and Fig. 7 Supplement 1A-C). The same measurements in control cells transfected with only Green cGull failed to demonstrate a clear dose-dependent response with respective increases of 17.5%, 39.8%, and 28.2% (Fig. 7B-D, F and Fig. 7 Supplement 1A-C). Area under the curve analysis confirmed significant differences between Gyc76C and control conditions (Fig. 7G). Exogenous protein expression levels were confirmed with post-hoc staining of Gyc76C-HA and Green cGull (Fig. 7 Supplement 1D). Several cells were found to express Green cGull independent of Gyc76C expression, explaining absence of strong response to ITPa in some cells. Finally, staining intensities were plotted against peak live-cell fluorescence increases, revealing weak negative correlations between exogenous protein expression and increases in cGMP signal (Fig. 7 Supplement 1D-F). The observed correlations may reflect lower dynamic ranges due to higher baseline fluorescence in cells receiving more exogenous DNA. Taken together, these live-imaging results strongly suggest that ITPa can induce increased cGMP production through Gyc76C.

**Figure 7:**
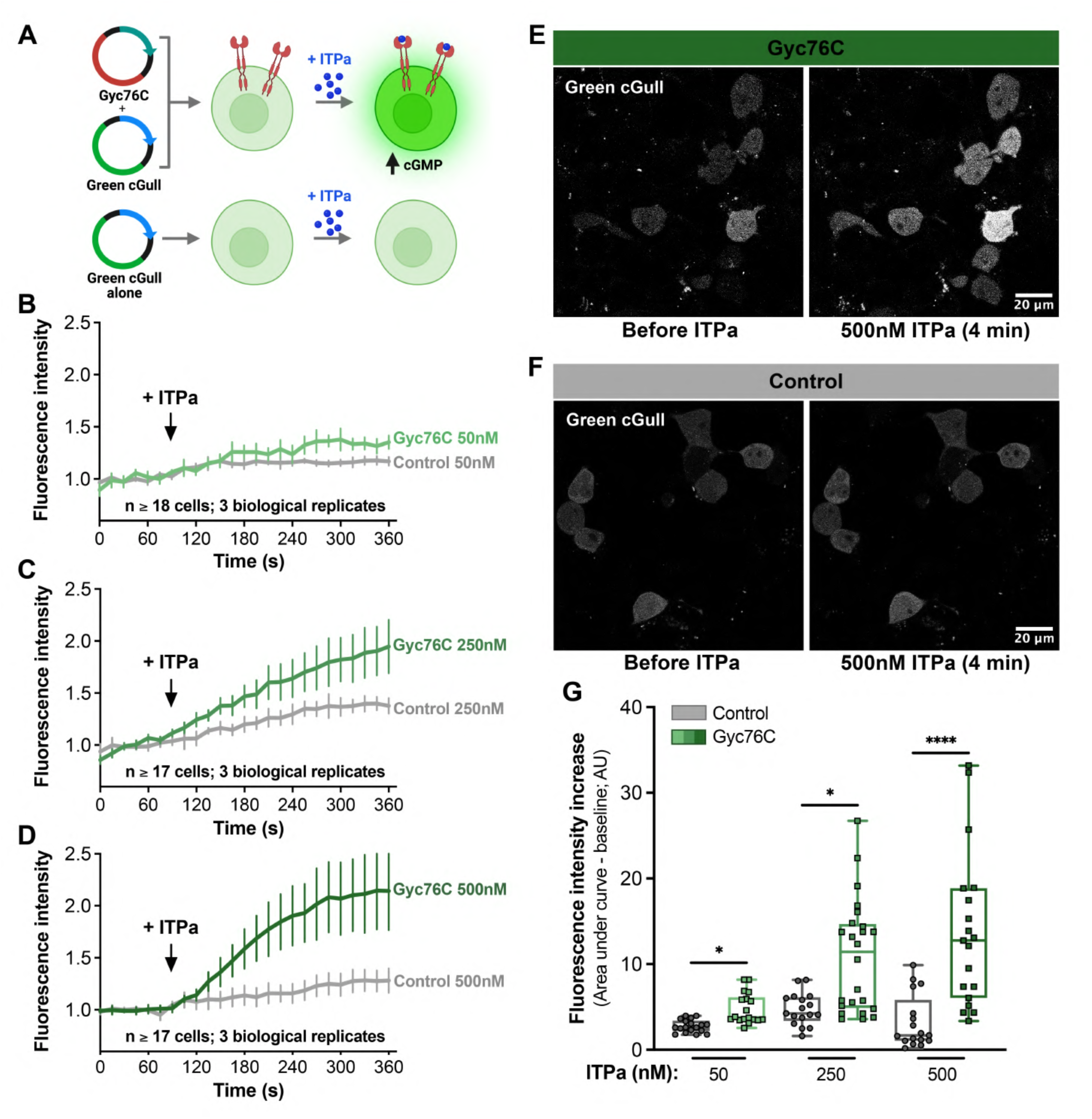
ITPa activates Gyc76C heterologously expressed in HEK293T cells. **(A)** Schematic of the heterologous assay used to functionally characterize Gyc76C. Created with BioRender.com. Application of **(B)** 50mM, **(C)** 250mM or **(D)** 500nM *Drosophila* ITPa to HEK293T cells transiently expressing Green cGull (cGMP sensor) and Gyc76C results in a dose-dependent increase in fluorescence compared to control cells which do not express Gyc76C. Graphs represent the mean fluorescence of 17-18 cells. Representative images showing fluorescence in **(E)** HEK293T cells expressing Gyc76C and **(F)** those without Gyc76C before and 4 mins after the addition of 500nm ITPa. **(G)** Area under the curve analysis demonstrates significant differences in Green cGull fluorescence increases between experimental and control conditions; * p < 0.05 and **** p < 0.0001 as assessed by nonparametric one-way ANOVA followed by Dunn’s test for multiple comparisons.

### ITPa-producing neurons are activated and release ITPa under desiccation

Having validated Gyc76C as a functional ITPa receptor, we next wanted to determine the context(s) during which ITP signaling is active *in vivo*. *ITP* expression has been shown to be upregulated during desiccation (Galikova *et al*., 2018). However, whether this increased transcription is also coupled with increased ITP signaling is unknown. Consistent with its role as an anti-diuretic hormone, we hypothesized that ITP signaling is increased under desiccation. To test this hypothesis, we employed CaLexA (Masuyama *et al*., 2012), a transcriptional reporter of neuronal activity, to monitor the activity of L-NSC^ITP^ in flies exposed to different contexts that challenge their osmotic homeostasis. In agreement with our prediction, L-NSC^ITP^ are more active (indicated by increased GFP immunofluorescence) in both males (Fig. 8A) and females (Fig. 8B) that were desiccated compared to flies that were kept under normal conditions. Moreover, L-NSC^ITP^ activity returned to normal levels in rehydrated flies that were previously exposed to dessication (Fig. 8A, B). In order to confirm that increased neuronal activity translates into increased peptide release, we independently quantified ITPa immunofluorescence in different subsets of ITPa-producing brain neurons (Fig. 8C, D). We observed reduced fluorescence in ITPa-expressing NSC as well as clock neurons in both males (Fig. 8C) and females (Fig. 8D) that had been exposed to desiccation stress. ITPa immunofluorescence returned to normal levels in desiccated flies that were then allowed to rehydrate. Since *ITP* mRNA is upregulated during desiccation, reduced immunofluorescence indicates increased release and not decreased peptide synthesis. Hence, not only do the L-NSC^ITP^ release ITPa into the circulation during desiccation, but the 5^th^-LN_v_ and LN_d_^ITP^ likely release ITPa within the brain to modulate other circuits during desiccation. Together, these results demonstrate that ITP neurons are activated and release ITPa during desiccation.

**Figure 8:**
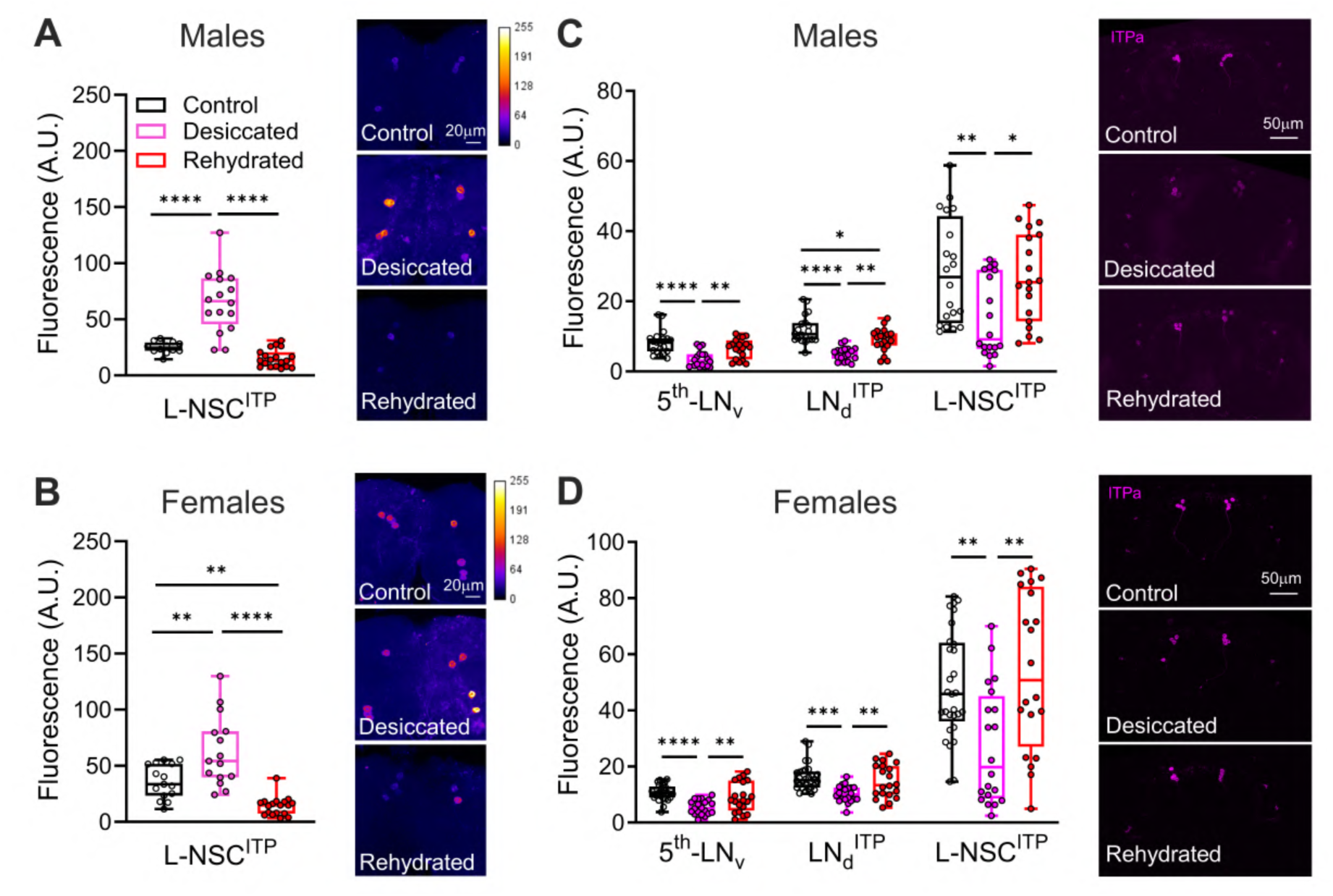
ITPa neurons are active and release ITPa during desiccation. GFP immunofluorescence, indicative of calcium levels and measured using the CaLexa reporter, is increased in L-NSC^ITP^ of **(A)** male and **(B)** female flies exposed to desiccation. The GFP intensity returns to control levels in flies that were rehydrated following desiccation. ITPa immunofluorescence, indicative of peptide levels, is lowered in 5^th^-LN_v_, LN_d_^ITP^ and L-NSC^ITP^ of **(C)** male and **(D)** female flies exposed to desiccation. ITPa peptide levels recover to control levels in flies that were rehydrated following desiccation. Lower peptide levels during desiccation indicates increased release. For all panels, * p < 0.05, ** p < 0.01, *** p < 0.001, **** p < 0.0001 as assessed by one-way ANOVA followed by Tukey’s multiple comparisons test.

### ITP knockdown in ITP-RC-T2A-GAL4 cells impacts osmotic and metabolic homeostasis

Insect ITP is evolutionarily related to CHH, which as its name indicates, regulates glucose homeostasis in crustaceans (Chen *et al*., 2020). A previous study employing ubiquitous ITP knockdown and overexpression suggests that *Drosophila* ITP also regulates feeding and metabolic homeostasis (Galikova and Klepsatel, 2022) in addition to osmotic homeostais (Galikova *et al*., 2018). However, given the nature of the genetic manipulations (ectopic ITPa overexpression and knockdown of *ITP* in all tissues) utilized in those studies, it is difficult to parse the effects of ITP signaling from ITPa-producing neurons. To fill this gap and understand the role of ITP signaling in regulating osmotic and metabolic homeostasis *in vivo*, we specifically knocked down *ITP* using the *ITP-RC-T2A-GAL4*, which includes all the ITPa-expressing neurons. To avoid any developmental effects, we combined temperature-sensitive tubulinGAL80 with the GAL4 line (here referred to as *ITP-RC-GAL4^TS^*) to restrict *ITP* knockdown specifically to the adult stage. For these experiments, both the *ITP-RNAi* and the control *luciferase RNAi* lines were first backcrossed for five generations into the wild-type background to minimize genetic background effects. We first successfully confirmed the effectiveness of *ITP-RNAi* by quantifying ITPa immunofluorescence in the brains of control and *ITP* knockdown flies (Fig. 9A). In agreement with the anti-diuretic effects of ITPa *ex vivo*, *ITP* knockdown resulted in reduced desiccation tolerance (Fig. 9B). This is likely a result of reduced water retention (Fig. 9C) and increased defecation (Fig. 9D). This reduced water content was also evident from the visibly shrunken abdomens of *ITP* knockdown females compared to controls (Fig. 9E).

**Figure 9:**
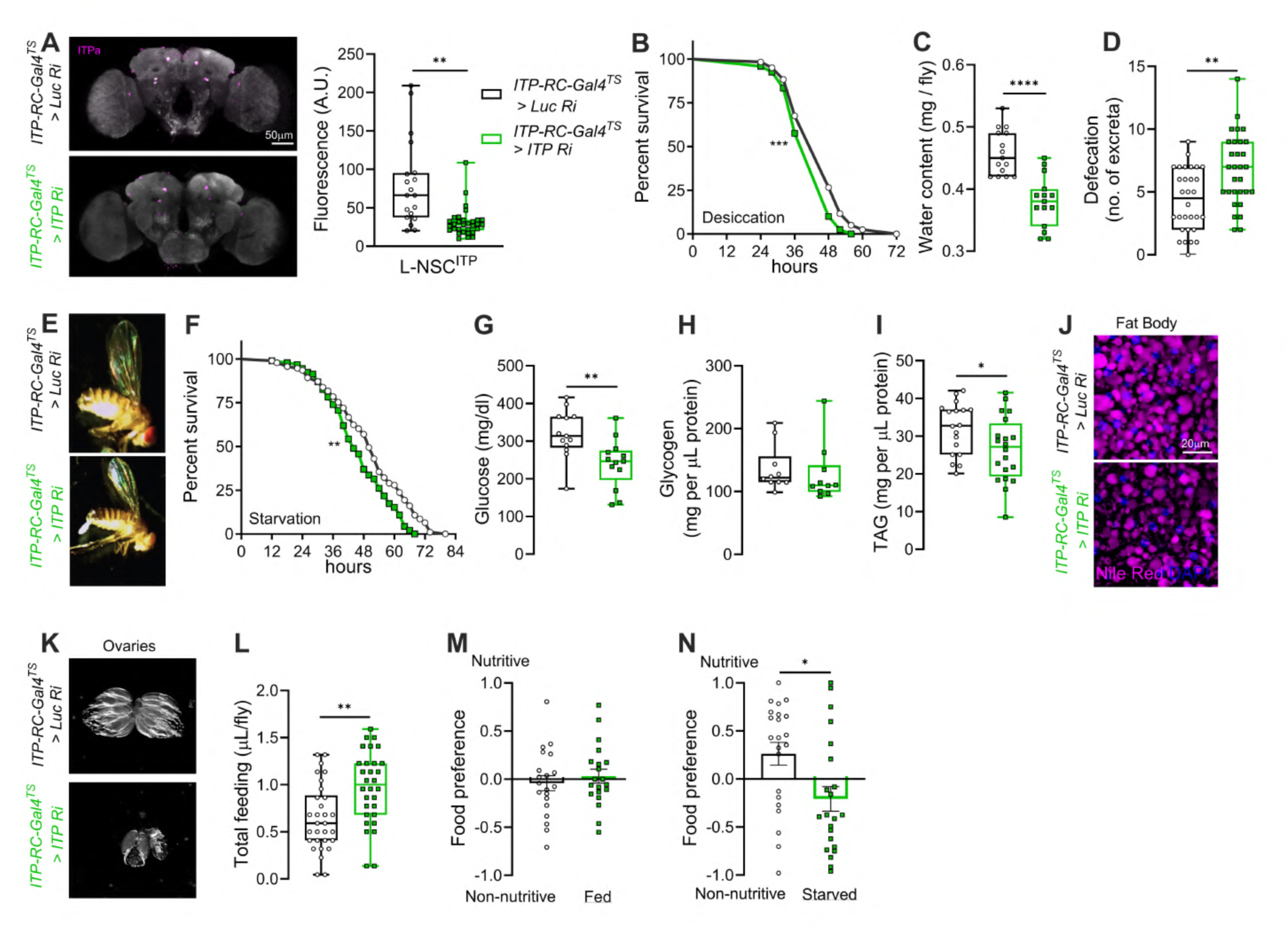
Knockdown of *ITP* in adult female *Drosophila* impacts metabolic homeostasis, feeding and associated behaviors. **(A)** ITPa immunofluorescence is reduced in the L-NSC^ITP^ neurons of flies in which *ITP* was knocked down using *ITP-RC-GAL4^TS^* (*ITP-RC-T2A-GAL4* combined with temperature-sensitive tubulin-GAL80). Flies with *ITP* knockdown are **(B)** less resistant to desiccation tolerance, **(C)** have reduced water content, **(D)** increased defecation, and **(E)** shrunken abdomen. *ITP* knockdown flies **(F)** survive less under starvation, **(G)** have lower levels of circulating glucose, and **(H)** unaffected glycogen levels. However, reduced ITP signaling results in **(I and J)** less lipid levels (TAG = triacylglyceride), and **(K)** smaller ovaries. Moreover, *ITP* knockdown flies exhibit **(L)** increased feeding (over 24 hours) and **(M and N)** defects in preference for nutritive sugars when starved for 18 hours prior to testing. Abbreviations: *Luc Ri, luciferase RNAi*; *ITP Ri, ITP RNAi*. For **B and F**, ** p < 0.01, *** p < 0.001, as assessed by Log-rank (Mantel-Cox) test. For all others, * p < 0.05, ** p < 0.01, and **** p < 0.0001 as assessed by unpaired *t* test.

Beyond its effects on osmotic balance, *ITP* knockdown in *ITP-RC* neurons also compromised metabolic homeostasis. Females with *ITP* knockdown exhibited reduced survival under starvation (Fig. 9F), prompting us to assess their circulating and stored macronutrients. As expected based on lowered starvation tolerance, these flies displayed significantly lower circulating glucose (Fig. 9G) and lipid stores in the fat body (Fig. 9I, J). Glycogen levels, however, remained unaltered compared to controls (Fig. 9H). As a consequence of these depleted energy reserves, the size of the ovaries was reduced in *ITP* knockdown females (Fig. 9K). Surprisingly, the reduction in energy reserves was not attributable to decreased food intake. On the contrary, *ITP* knockdown flies consumed more food than controls (Fig. 9L). Independently, we also assayed the food preference of flies when given a choice between a nutritive sugar and a sweeter non-nutritive sugar since it can report deficits in mechanisms that monitor internal metabolic state. While fed flies of both the control and experimental genotypes showed no preference (Fig. 9M), starved control flies exhibited a shift towards caloric nutritive sugars (Fig. 9N), which reflects their drive to restore energy balance. In contrast, starved *ITP* knockdown flies did not display this shift in preference towards nutritive sugars (Fig. 9N), indicating a disruption in the integration of internal metabolic cues with taste-driven food selection.

Collectively, these results demonstrate that disrupting ITP signaling from *ITP-RC* neurons leads to profound systemic effects on osmotic regulation, metabolic balance and feeding behaviors, underscoring the pivotal role of ITP in coordinating diverse physiology and behaviors.

### ITPa overexpression in ITP-RC-T2A-GAL4 cells modulates osmotic and metabolic homeostasis

The *ITP-RNAi* used here targets all *ITP* isoforms. Hence, the phenotypes observed following *ITP* knockdown cannot directly be attributed to ITPa. Therefore, we complemented these analyses by specifically overexpressing *ITPa* in adult females using *ITP-RC-GAL4^TS^* (Fig. 10). We first confirmed that driving *UAS-ITPa* with *ITP-RC-GAL4^TS^* indeed results in increased ITPa peptide levels in L-NSC^ITP^ (Fig. 10A). In agreement with our *ex vivo* secretion data, ITPa overexpression improves desiccation tolerance (Fig. 10B), likely due to increased water retention, since flies overexpressing ITPa had higher body water content (Fig. 10C) and bloated abdomens (Fig. 10D). Independently, we also assessed recovery from chill-coma as an indirect measure of flies’ ionoregulatory capacity (MacMillan *et al*., 2012). ITPa overexpression had no impact on chill coma recovery (Fig. 10 Supplement 1A) and tolerance to salt stress (Fig. 10 Supplement 1B). With regard to metabolic physiology, flies with ITPa overexpression survive longer under starvation (Fig. 10E). These flies had reduced glucose levels (Fig. 10F) but their glycogen levels were unaltered (Fig. 10G). ITPa overexpression also led to increased lipid levels (Fig. 10H). In addition, flies with ITPa overexpression showed defects in preference between nutritive versus non-nutritive sugar (Fig. 10I, J) and had larger ovaries (Fig. 10K). Thus, ITPa overexpression largely results in opposite phenotypes compared to those seen following *ITP* knockdown.

**Figure 10:**
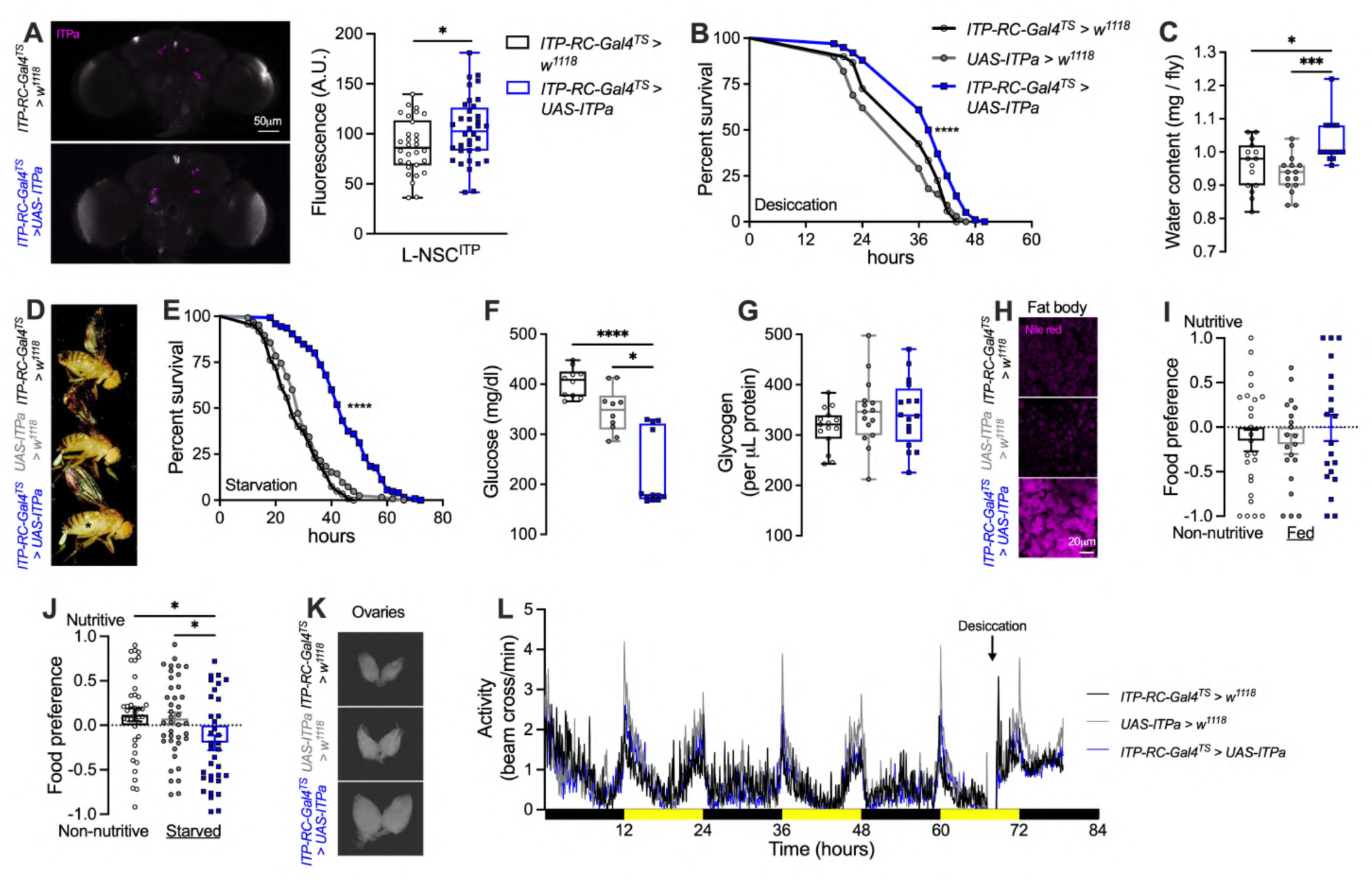
ITPa overexpression in adult female *Drosophila* impacts osmotic and metabolic homeostasis, feeding and related behaviors. **(A)** Overexpression of *ITPa* using *ITP-RC-GAL4^TS^* results in increased ITPa immunofluorescence in L-NSC^ITP^. ITPa overexpression results in **(B)** increased desiccation tolerance, **(C)** increased water content and **(D)** a slightly bloated abdomen (marked by an asterisk). ITPa overexpression causes **(E)** increased starvation tolerance, **(F)** reduced circulating glucose levels but has no effect on **(G)** glycogen levels. **(H)** The size of neutral lipid droplets (stained with Nile red) is increased in flies with ITPa overexpression. **(I and J)** These flies also exhibit defects in preference for nutritive sugars when starved for 16 hours prior to testing. **(K)** ITPa overexpression flies have enlarged ovaries. **(L)** ITPa overexpression has no effect on locomotor activity under fed or desiccating conditions. Black bars indicate night-time and yellow bars indicate daytime. All experiments were performed at 29°C. For **B and E**, **** p < 0.0001, as assessed by Log-rank (Mantel-Cox) test. For **A**, * p < 0.05 as assessed by unpaired *t* test. For all other experiments, * p < 0.05, *** p < 0.001, **** p < 0.0001 as assessed by one-way ANOVA followed by Tukey’s multiple comparisons test. For clarity, significant pairwise differences compared to only the experimental treatment are indicated.

Independently, since ITPa is released from both clock neurons and L-NSC^ITP^ during desiccation, we asked whether ITPa overexpression in both neuron types affected rhythmmic locomotor activity under normal and desiccating conditions. We did not observe any drastic differences in locomotor activity of flies kept under normal conditions and subsequently transferred to empty vials (desiccation conditions) (Fig. 10L). Therefore, the function of ITPa released from clock neurons during desiccation remains to be determined.

### ITPa signals via Gyc76C in the renal tubules to modulate osmotic homeostasis

We next explored the role of ITP signaling via Gyc76C in maintaining osmotic homeostasis *in vivo*. In order to test if the effects on osmotic homeostasis were mediated via Gyc76C in the renal tubules, we monitored osmotic and ionic/salt stress tolerance of flies in which Gyc76C was specifically knocked down in MT principal or stellate cells using the *uro-GAL4* and *c724-GAL4*, respectively (Fig. 11). As expected, flies with Gyc76C knockdown in the MTs exhibit reduced tolerance to desiccation irrespective of the cell type, principal or stellate, being targeted (Fig. 11A, D). Interestingly, salt stress impacted flies differently depending on the cell type in which Gyc76C was knocked down. Principal cell knockdown led to increased survival (Fig. 11B), whereas stellate cell knockdown resulted in reduced survival (Fig. 11E), which could reflect functional differences between the two cell types. Lastly, Gyc76C knockdown in either MT cell type increased the time taken to recover from chill-coma, highlighting deficits in the ability to maintain ionic homeostasis (Fig. 11C, F). In summary, ITPa is released into the circulation during desiccation and modulates the MTs to promote tolerance to osmotic and ionic stresses. This evidence suggests that the hormonal effect of ITPa is likely mediated via Gyc76C expressed in the stellate and principal cells of the MTs.

**Figure 11:**
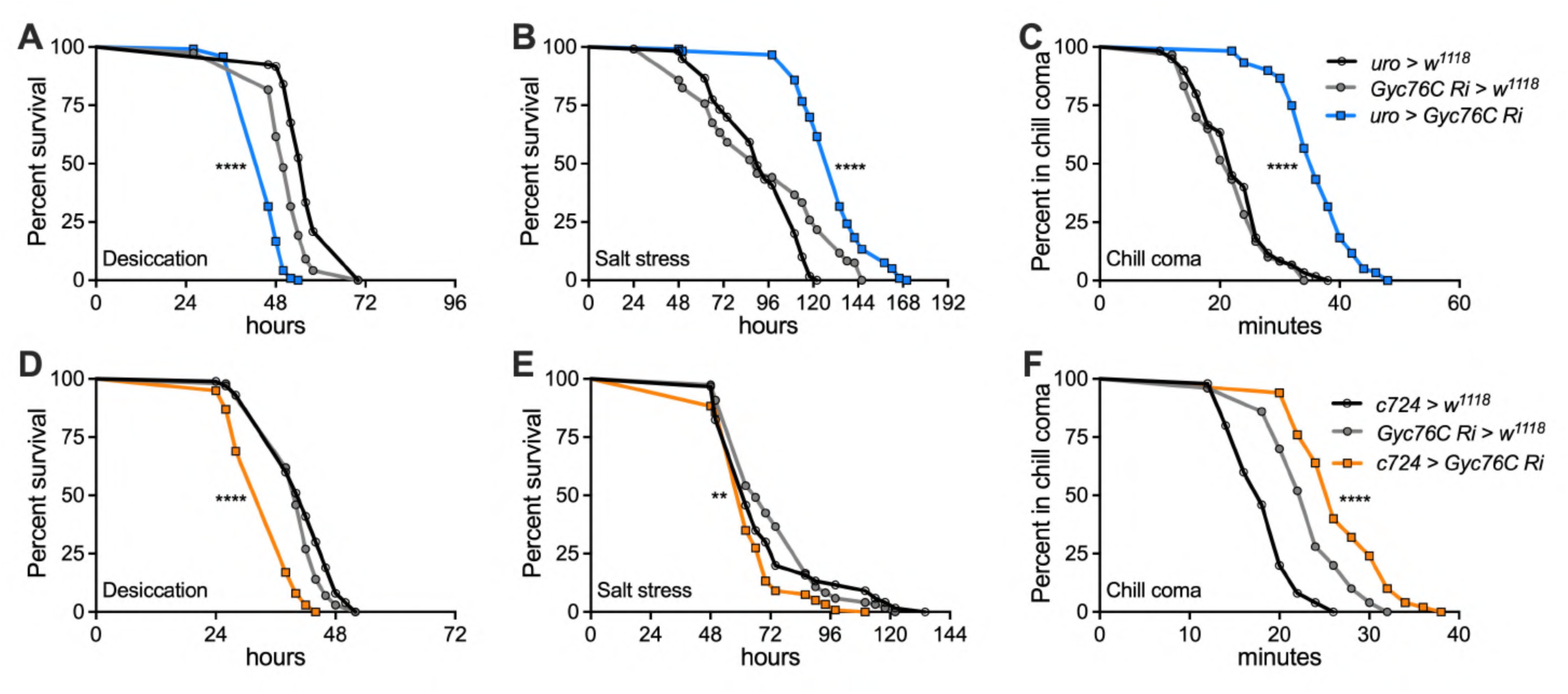
Female Malpighian tubule specific knockdown of *Gyc76C* impacts osmotic and ionic homeostasis. Knockdown of *Gyc76C* in both the **(A)** principal cells of renal tubules using *uro-GAL4* and **(D)** stellate cells using *c724-GAL4* reduces desiccation tolerance. *Gyc76C* knockdown in **(B)** principal cells increases survival under salt stress whereas knockdown in **(E)** stellate cells lowers survival. **(C and F)** *Gyc76C* knockdown in principal or stellate cells increases the time taken for recovery from chill-coma. Abbreviation: *Gyc76C Ri, Gyc76C RNAi.* For all panels, ** p < 0.01, **** p < 0.0001, as assessed by Log-rank (Mantel-Cox) test.

### ITPa-Gyc76C signaling to the fat body influences metabolic physiology and associated behaviors

Having identified the inter-organ pathway via which ITPa modulates osmotic homeostasis, we next wanted to characterize the pathway regulating metabolic physiology. Since Gyc76C is expressed in the fat body and only the female IPCs, but not in AKH-producing cells, we hypothesized that ITP primarily regulates metabolic homeostasis via direct signaling to the fat body. To test this prediction, we specifically knocked down *Gyc76C* in the female fat body using *yolk-GAL4*. Flies with *Gyc76C* knockdown in the fat body exhibit a drastic reduction in starvation tolerance compared to control flies (Fig. 12A). Remarkably, these flies start dying after only 4 hours of starvation, whereas control flies can normally tolerate at least 24 hours of starvation. Therefore we investigated whether Gyc76C signaling to the fat body impacts energy stores. In agreement with reduced starvation survival, *Gyc76C* knockdown flies have lower hemolymph glucose (Fig. 12B), unaltered glycogen levels (Fig. 12C), and lower lipids in the fat body (Fig. 12D, E) compared to controls. Hence, lower lipid levels, especially in the fat body, likely contribute to reduced starvation survival. Interestingly, the reduction in energy stores is not due to decreased food intake because *Gyc76C* knockdown flies fed more than controls (Fig. 12F). In addition, these flies displayed altered food preferences. Specifically, they prefered yeast over sucrose (Fig. 12G), possibly to mitigate protein or general caloric deficits since the protein in yeast yields greater caloric value than sugar. Independently, we also assayed the preference of flies for nutritive versus non-nutritive sugars. While there was no preference for nutritive or non-nutritive sugar in fed flies (Fig. 12H), control flies showed increased preference for nutritive sugar during starvation (Fig. 12I). Conversely, starved *Gyc76C* knockdown flies displayed a slight preference for non-nutritive sugar over nutritive sugar (Fig. 12I), suggesting disrupted integration of taste signals with the internal metabolic state. In summary, these experiments indicate that Gyc76C signaling in the fat body is vital in regulating feeding, metabolic homeostasis and consequently survival.

**Figure 12:**
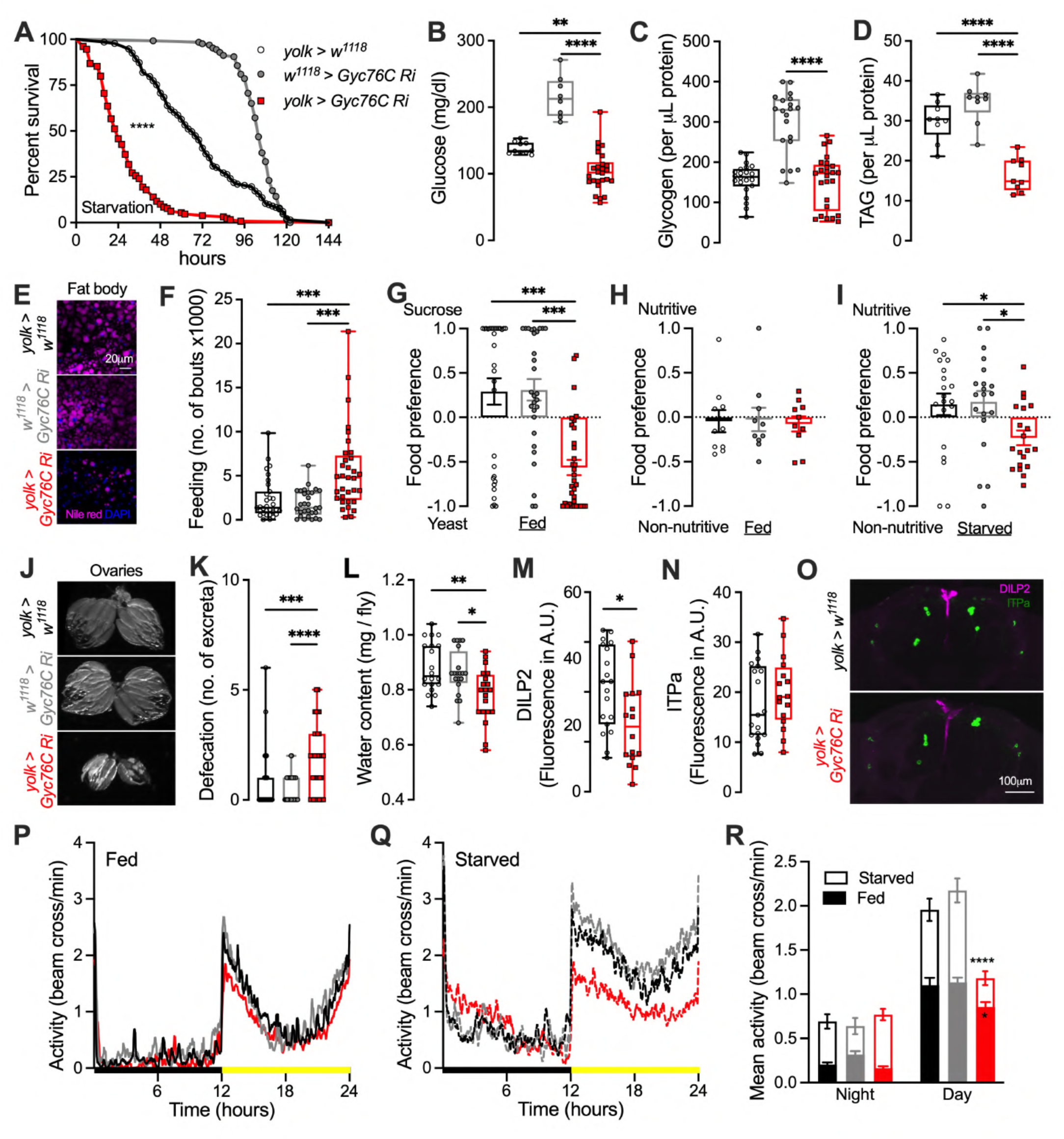
*Gyc76C* knockdown in the female fat body using *yolk-GAL4* impacts metabolic homeostasis, feeding and associated behaviors. Flies with fat body specific *Gyc76C* knockdown with *UAS-Gyc76C RNAi* (*#106525*) are **(A)** extremely susceptible to starvation and **(B)** have reduced glucose levels. **(C)** Glycogen levels are unaltered in flies with fat body specific *Gyc76C* knockdown. **(D and E)** However, lipid levels (TAG = triacylglyceride) are drastically reduced. *Gyc76C* knockdown flies exhibit **(F)** increased feeding (over 24 hours), **(G)** a preference for yeast over sucrose, and **(H and I)** defects in preference for nutritive sugars when starved for 4 hours prior to testing. Flies with *Gyc76C* knockdown in the fat body have **(J)** smaller ovaries, they **(K)** defecate more and have **(L)** reduced water content than the controls. For **K,** number of excreta counted over 2 hours. *Gyc76C* knockdown also impacts **(M)** DILP2 peptide levels **(N)** but not ITPa levels in the neurosecretory cells. CTCF = Corrected Total Cell Fluorescence. **(O)** Representative confocal stacks showing DILP2 and ITPa immunostaining. *Gyc76C* knockdown flies also display reduced daytime locomotor activity under **(P)** fed and **(Q)** and starved conditions compared to controls. Black bars indicate night-time and yellow bars indicate daytime. **(R)** Average night and daytime activity over one day under fed and starved conditions. For **A**, **** p < 0.0001, as assessed by Log-rank (Mantel-Cox) test. For **M and N,** * p < 0.05 as assessed by unpaired *t* test. For all others, * p < 0.05, ** p < 0.01, *** p < 0.001, **** p < 0.0001 as assessed by one-way ANOVA followed by Tukey’s multiple comparisons test. For clarity, significant pairwise differences compared to only the experimental treatment are indicated.

Next, we examined if fat body specific *Gyc76C* knockdown impacts other tissues and behaviors. We first observed that *Gyc76C* knockdown flies had drastically shrunken ovaries (Fig. 12J). In addition, knockdown flies also defecated more than controls (Fig. 12K) and relatedly had a lower water content (Fig. 12L). These effects could either be caused by altered feeding and metabolism and/or via an indirect impact on insulin and ITP signaling amongst other pathways. Particularly, reduced insulin and ITP signaling could result in the observed reproductive and excretory phenotypes, respectively. Therefore, we quantified DILP2 and ITPa peptide levels in the brain NSC following knockdown of *Gyc76C* in the fat body. Indeed, knockdown flies have reduced DILP2 peptide levels (Fig. 12M, O). However, we did not observe any differences in ITPa peptide levels in L-NSC^ITP^ (Fig. 12N, O). These results suggest that the shrunken ovaries could be directly caused by reduced nutrient stores as well as via an indirect effect on insulin and possibly juvenile hormone signaling since L-NSC^DH31^ innervate the corpora allata. The increased defecation, on the other hand, could likely be a consequence of increased feeding or due to an impact on another osmoregulatory pathway. To further substantiate the fat body specific role of Gyc76C, we repeated several of the aforementioned analyses using an independent RNAi line (*Gyc76C RNAi #2*) and observed comparable phenotypes. Flies with *Gyc76C* knockdown in the fat body using *Gyc76C RNAi #2* exhibited reduced tolerance to both desiccation (Fig. 12 Supplement 1A) and starvation stress (Fig. 12 Supplement 1B). These flies also displayed reduced lipid stores (Fig. 12 Supplement 1C) and smaller ovaries (Fig. 12 Supplement 1D). Our findings provide strong evidence that disruption of Gyc76C signaling in the fat body exerts profound systemic effects on multiple aspects of physiology and behavior.

Lastly, we monitored general locomotor activity and starvation induced hyperactivity, the latter of which is largely governed by AKH, insulin and octopamine signaling (Lee and Park, 2004, Yu *et al*., 2016b, Pauls *et al*., 2021). Flies with *Gyc76C* knockdown in the fat body displayed reduced daytime activity when kept under either fed or starved conditions for one day (Fig. 12P-R). Hence, Gyc76C signaling in the fat body does not appear to impact starvation-induced hyperactivity. However, the effect on general locomotor activity led us to examine the activity of fed flies in more detail over a longer time course. For this, we monitored the activity of flies for 10 days under 12:12-hour light/dark cycles and a subsequent 10 days under constant darkness (Fig. 12 Supplement 2A, B). While *Gyc76C* knockdown flies displayed reduced activity on day 1, the average activity of these flies over days 2 to 6 was not significantly different from the controls (Fig. 12 Supplement 2C, D). Interestingly, flies with *Gyc76C* knockdown in the fat body appeared to be more sensitive to differences in light cues, showing a strong reduction in locomotor activity only when switched to constant darkness from 12:12-hour light dark cycles (Fig. 12 Supplement 2A, B).

In conclusion, the phenotypes seen following *Gyc76C* knockdown in the fat body largely mirror those seen following *ITP* knockdown in *ITP-RC* neurons, providing further support that ITPa mediates its effects via Gyc76C.

### Synaptic and peptidergic connectivity of ITP neurons

After characterizing the functions of ITP signaling to the renal tubules and the fat body, we wanted to identify pathways regulating ITP signaling and its downstream neuronal targets. To address this, we took advantage of the recently completed FlyWire adult brain connectome (Dorkenwald *et al*., 2024, Schlegel *et al*., 2024) to identify pre- and post-synaptic partners of ITP neurons. ITP neurons have a characteristic morphology which was used to identify them in the connectome (Fig. 13A) (McKim *et al*., 2024, Reinhard *et al*., 2024). LN_d_ ^ITP^ and 5^th^-LN_v_ displayed numerous input and output synapses in the brain (Fig. 13B and Figure 13 Supplement 1). In particular, these neurons have extensive synaptic output in the superior lateral protocerebrum where Gyc76C-expressing dorsal clock neurons reside (Fig. 5M and Fig. 5 Supplement 2F). Consistent with the high number of synapses, both LN_d_ ^ITP^ and 5^th^-LN_v_ receive inputs and provide outputs to a broad range of neurons (Fig. 13C, D). In addition, both cell types are upstream of at least one pair of DH_31_-expressing NSC (L-NSC^DH31^) (Fig. 13D). However, it is not yet clear whether these NSC are the same ones as the L-NSC^DH31^ that co-express ITPa (Fig. 1B, E), since there are three pairs of L-NSC^DH31^ in the adult brain (Reinhard *et al*., 2024). Therefore, we did not examine the synaptic connectivity of L-NSC^DH31^ here. Interestingly, LN_d_ ^ITP^ receive indirect inputs from VP1l thermo/hygrosensory neurons (Fig. 13E) (Choi *et al*., 2022). LN_d_ ^ITP^ coexpress neuropeptide F (NPF) and could be the same NPF-expressing neurons which have recently been implicated in water seeking during thirst (Ramirez *et al*., 2025). This connectivity could also explain why LN_d_ ^ITP^ show reduced ITPa immunolabelling (indicating ITPa release) following desiccation (Fig. 8C, D).

**Figure 13:**
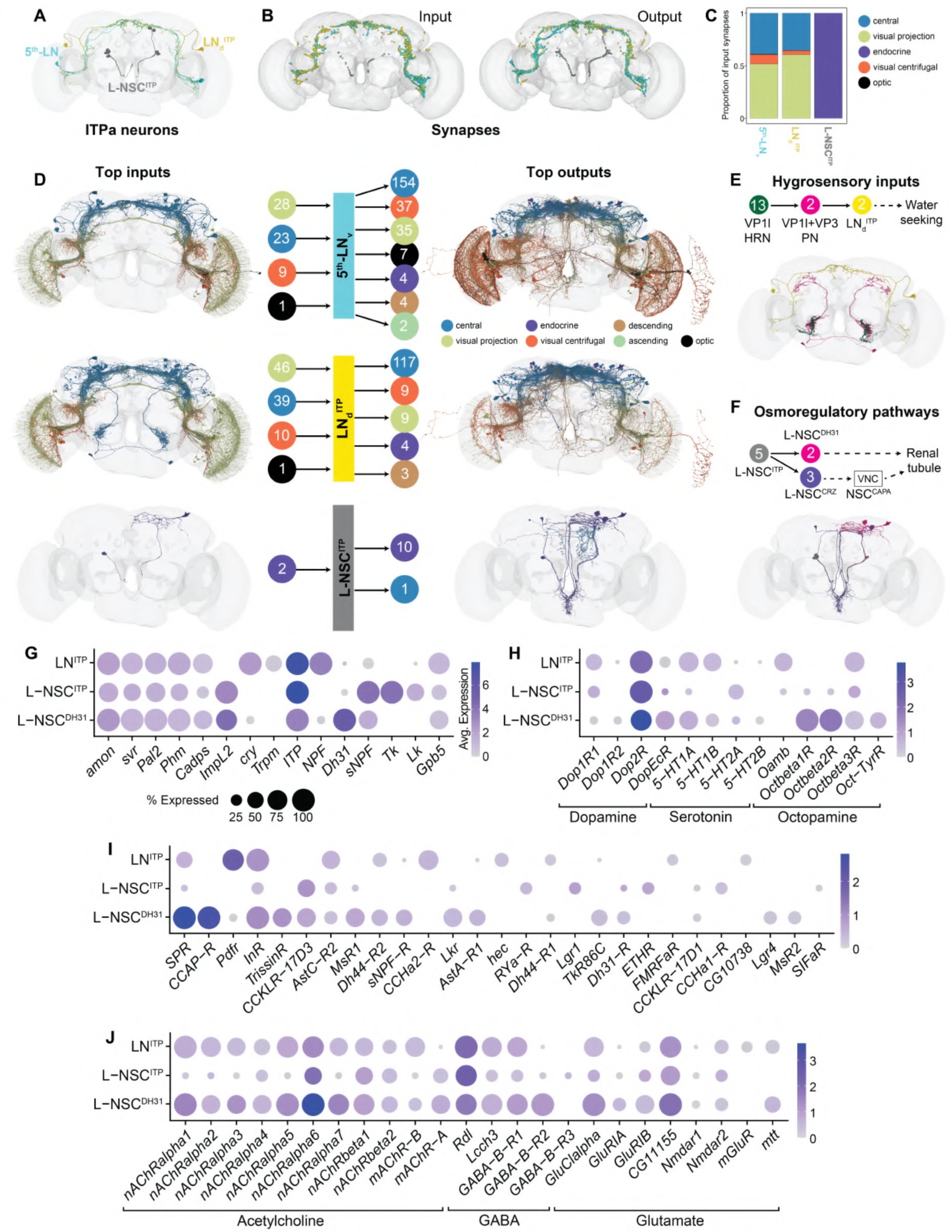
Inputs and outputs of ITP neurons based on connectomics and single-cell transcriptomics. **(A)** Reconstruction of ITPa-expressing neurons using the complete electron microscopy volume of the adult female brain (data retrieved from the FlyWire platform). Four pairs of lateral neurosecretory cells (L-NSC^ITP^) are grey, 5^th^ ventrolateral neurons (5^th^-LN_v_) are cyan, and dorsolateral neurons (LN_d_^ITP^) are yellow. Diuretic hormone 31 (DH_31_)-expressing lateral neurosecretory cells (L-NSC^DH31^) are not shown since it is unclear which of the three pairs of L-NSC^DH31^ co-expresses ITPa. **(B)** Location of input and output synapses are colored according to the ITP neuron type. **(C)** Proportion of input synapses (grouped by super class annotations for the FlyWire connectome (Schlegel *et al*., 2024)) to each ITP neuron type. **(D)** Reconstructions of neurons from different super classes providing inputs to (left) and receiving outputs from (right) 5^th^-LN_v_, LN_d_^ITP^ and L-NSC^ITP^. Only the top 10 cell types are shown here. (Middle) Number of neurons, categorized by super class, providing inputs to and receiving outputs from 5^th^-LN_v_, LN_d_^ITP^ and L-NSC^ITP^. **(E)** Thermo/hygrosensory input pathway to LN_d_^ITP^. **(F)** Output from L-NSC^ITP^ to other osmoregulatory hormone-producing cells. **(G)** Identification of single-cell transcriptomes representing different subsets of ITPa-expressing neurons in the adult brain dataset (Davie *et al*., 2018). Since both the 5^th^-LN_v_ and LN_d_^ITP^ co-express *ITP*, *cryptochrome* (*cry*) and *neuropeptide F* (*NPF*), these cells are grouped as LN^ITP^. All three sets of neurons express genes required for neuropeptide processing and release (*amon*, *svr*, *Pal2*, *Phm* and *Cadps*) and were identified based on the neuropeptides (*ITP*, *NPF, Dh31, sNPF* and *Tk*) they express. Dot plots showing expression of **(H)** monoamine, **(I)** neuropeptide and **(J)** neurotransmitter receptors in different sets of ITPa neurons.

In contrast to LN ^ITP^ and 5^th^-LN clock neurons, L-NSC^ITP^ form few significant synaptic connections within this brain volume (Fig. 13B, D and Figure 13 Supplement 1). Since these neurons are neurosecretory in nature, their peptides are released from axon terminations in neurohemal areas outside the brain, where regulatory inputs could be located (Dircksen *et al*., 2008, Kahsai *et al*., 2010, McKim *et al*., 2024). It is worth noting that L-NSC^ITP^ output onto other NSC subtypes which secrete osmoregulatory hormones like DH_31_ and corazonin (CRZ) (Fig. 13F). DH_31_ is a diuretic hormone whereas CRZ can inhibit CAPA, another diuretic hormone (Zandawala *et al*., 2021).

Since L-NSC^ITP^ receive few synaptic inputs, we hypothesized that their activity, especially during desiccation, is regulated either by cell autonomous osmosensing or by paracrine and hormonal modulators which transmit the signal from other central or peripheral osmosensors. To address this, we mined single-cell transcriptomes of different subsets of *ITP*-expressing neurons (Fig. 13G and Fig. 13 Supplement 2A) from whole brain and VNC datasets (Davie *et al*., 2018, Allen *et al*., 2020) based on markers identified here and previously (Kahsai *et al*., 2010, Reinhard *et al*., 2024). To assess if *ITP*-expressing neurons are cell autonomously osmosensitive, we first examined the expression of transient receptor potential (TRP) and pickpocket (ppk) channels which have been shown to confer osmosensitivity to cells (Sharif-Naeini *et al*., 2008, Cameron *et al*., 2010). Although *Trpm*, a TRP channel, was expressed in LN^ITP^ (Fig. 13G), we did not detect expression of any TRP or ppk channels in L-NSC^ITP^ and L-NSC^DH31^ (not shown). Thus, unless other (non-charaterized) osmosensors are expressed in L-NSC^ITP^, the internal state of thirst/desiccation is likely conveyed to L-NSC^ITP^ via neuromodulators. Intriguingly, the dopamine receptor, *Dop2R*, is highly expressed in all ITP neuron subtypes (Fig. 13H and Fig. 13 Supplement 2B). Compared to the *ITP-*expressing neurons in the VNC, the brain neurons seem to be extensively modulated by different neuropeptides (Fig. 13I and Fig. 13 Supplement 2C), including those that regulate osmotic homeostasis (diuretic hormone 44, LK and DH_31_), and feeding and metabolic homeostasis (insulin, Ast-A, CCAP, drosulfakinin and sNPF). Lastly, with the exception of L-NSC^ITP^, all *ITP* neurons express high levels of neurotransmitter receptors (Fig. 13J and Fig. 13 Supplement 2D). This is consistent with fewer synaptic inputs to L-NSC^ITP^. In conclusion, the ITP neuron connectomes and transcriptomes provide the basis to functionally characterize signaling pathways regulating ITP signaling in *Drosophila*.

## Discussion

Insect ITPs are members of the multifunctional family of CHH/MIH neuropeptides that have been intensely investigated in crustaceans for their role in development, reproduction, and metabolism (Webster *et al*., 2012). In *Drosophila,* the three ITP isoforms (ITPa, ITPL1 and ITPL2) were until very recently among the very few neuropeptides whose receptors had not been identified. However, recently an ITPL2-activated GPCR, TkR99D, was identified (Xu *et al*., 2023) similar to the ITPL-activated BNGR-A24 in the moth *Bombyx* (Nagai *et al*., 2014). Thus, receptors for *Drosophila* ITPa and ITPL1 remained to be identified. Furthermore, the neuronal pathways and functional roles of the three ITP isoforms have remained relatively uncharted. Here, using a multipronged approach consisting of anatomical mapping, single-cell transcriptomics, *in vitro* tests of recombinant ITPa and genetic experiments *in vivo,* we comprehensively mapped the tissue expression of all three *ITP* isoforms and revealed roles of ITP signaling in regulation of osmotic and metabolic homeostasis via action on Malpighian (renal) tubules and fat body, respectively. We furthermore identified and functionally characterized a receptor for the amidated isoform ITPa, namely the mGC Gyc76C and analyzed its tissue distribution and role in systemic homeostasis. Lastly, we performed connectomics and single-cell transcriptomic analyses to identify synaptic and paracrine pathways upstream and downstream of ITP-expressing neurons. Together, our systematic characterization of ITP signaling establishes a tractable system to decipher how a small set of neurons integrates diverse inputs and orchestrates systemic homeostasis in *Drosophila* (Fig. 14).

**Figure 14:**
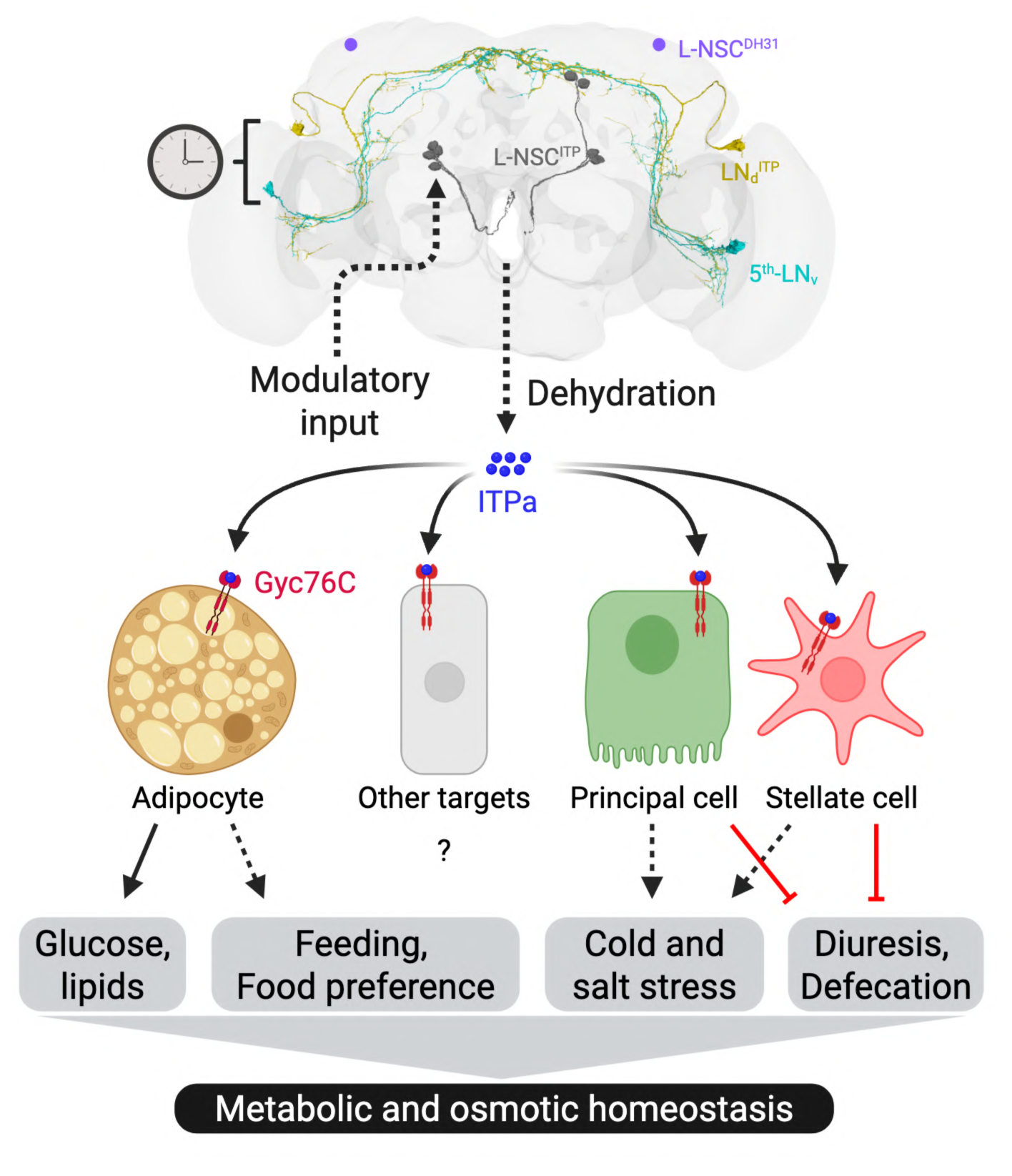
A schematic depicting ITP signaling pathways modulating metabolic and osmotic homeostasis in *Drosophila*. Different subsets of ITP neurons in the brain have been color-coded. LN_d_^ITP^ and 5^th^-LN_v_ are part of the circadian clock network and regulate clock-associated behaviors and physiology. L-NSC^ITP^ release ITPa into the circulation following dehydration and information regarding this internal state is likely conveyed to L-NSC^ITP^ by other neuromodulators. Following its release into the hemolymph, ITPa activates a membrane guanylate cyclase receptor Gyc76C on the adipocytes in the fat body, principal and stellate cells in the renal tubules, as well as other targets. These signaling pathways affect diverse behaviors and physiology to modulate metabolic and osmotic homeostasis. Dashed arrows depict pathways that remain to be clarified, solid arrows represent direct effects, and red bars represent inhibition. Created with BioRender.com.

### ITP neurons release multiple neuropeptides to regulate systemic homeostasis

ITPa action on renal tubules and fat body is very likely to be hormonal via the circulation since these tissues are not innervated by neurons. While there are peripheral cells that could possibly release ITPa into the circulation (Fig. 3J), we consider the eight L-NSC^ITP^ to be the major source of hormonal ITPa (and ITPL forms). This is based on the fact that these cells have large cell bodies, numerous dense core vesicles and extensive axon terminations for production and storage of large amounts of peptide. The smaller L-NSC^DH31^ have low levels of ITPa and are likely more suited to locally modulate the corpora allata and/or axon terminations of other *bona fide* NSC in that region. It is noteworthy that L-NSC^ITP^ and L-NSC^DH31^ express at least five neuropeptides each, with ITPa, ITPL1, sNPF and glycoprotein hormone beta 5 (Gpb5) being common across both cell types. Additionally, L-NSC^ITP^ express ITPL2, LK and TK, while L-NSC^DH31^ express DH_31_. While the functions of *Drosophila* ITPL1 are still unknown, the other neuropeptides have been shown to regulate osmotic and metabolic stress responses (Johnson *et al*., 2005, Kahsai *et al*., 2010, Zandawala *et al*., 2018a, Diaz-de-la-Pena *et al*., 2020). Interestingly, both L-NSC^ITP^ and L-NSC^DH31^ also express ImpL2, an insulin-binding protein which enables cells to sequester insulin (Bader *et al*., 2013, Galikova *et al*., 2018, Ghosh *et al*., 2022). These ITP neurons can thus act as a reservoir for insulin-like peptides and could release them along with other neuropeptides to modulate both osmotic and metabolic homeostasis. If we account for DILP2 (Bader *et al*., 2013), L-NSC^ITP^ can release up to eight neuropeptides, the most detected for a neuron type so far in *Drosophila.* Hence, understanding the mechanisms by which these cells are regulated can provide novel insights into their orchestrating actions in mediating systemic homeostasis.

Although we consider the L-NSC^ITP^ as the main players in hormonal release of ITP isoforms, other ITP producing neurons could also regulate peripheral tissues. For instance, iag neurons in the abdominal ganglia, which directly innervate the hindgut and rectum, likely modulate gut physiology. These neurons also produce multiple neuropeptides in addition to ITPa, namely ITPL1, CCAP, Ast-A and Gpb5. CCAP and Ast-A peptides could modulate hindgut contractility (Vanderveken and O’Donnell, 2014, Hillyer, 2018), whereas ITPa and Gpb5 could regulate water and ion reabsorption (Sellami *et al*., 2011). The concerted action of all these neuropeptides on the hindgut could thus facilitate osmotic homeostasis.

### When is ITPa released and how are ITP neurons regulated?

The release of ITPa from ITPa-expressing neurons in the brain appears to be regulated by the state of water and ion balance in the fly, as seen in our experiments measuring ITP neuron activity and ITPa peptide levels in desiccated and rehydrated flies. But how is this internal state of thirst/desiccation conveyed to ITP neurons? In mammals, osmotic homeostasis is regulated by vasopressin neurons in the hypothalamus (Voisin and Bourque, 2002). These neurons monitor changes in the osmotic pressure via intrinsic mechanosensitive channels. In addition, synaptic and paracrine inputs also regulate vasopressin release. Although the vasopressin signaling system has been lost in *Drosophila* (Nässel and Zandawala, 2019), other osmoregulatory systems such as ITP could have evolved similar mechanisms to monitor and consequently regulate the osmotic state of the animal. Our connectomic and single-cell transcriptome analysis indicates that information regarding the osmotic state is likely conveyed to ITP NSC indirectly via one or more neuromodulators released from other osmosensors. Since several receptors for neuromodulators are expressed in L-NSC^ITP^, it is difficult to predict which neuromodulators convey the thirst signal to ITP neurons. Nonetheless, it is tempting to speculate that this signal could be dopamine since *Dop2R* is highly expressed in ITP neurons (Fig. 13H) and dopaminergic neurons also track changes in hydration in mice (Grove *et al*., 2022). Future investigations are needed to explore the modulation of ITP neurons by dopamine and other modulators. Interestingly, the two pairs of clock neurons, 5^th^-LN_v_ and LN_d_^ITP^, also release ITPa during desiccation (Fig. 8C, D). The behavioral effects of ITPa signaling by clock neurons during desiccation remains to be discovered, since locomotor activity under normal and desiccation conditions was not affected following ITPa overexpression.

### Additional targets of ITPa-Gyc76C signaling

Gyc76C was identified in *Drosophila* as a mGC in the mid 1990s (Liu *et al*., 1995, McNeil *et al*., 1995), and has since been shown to play diverse roles in embryonic development of different epithelia (including renal tubules) and muscle (Patel *et al*., 2012, Patel and Myat, 2013, Schleede and Blair, 2015, Myat and Patel, 2016), axonal growth and guidance (Ayoob *et al*., 2004, Chak and Kolodkin, 2014), innate immunity (Iwashita *et al*., 2020) and salt stress tolerance (Overend *et al*., 2012). Given the crucial role of Gyc76C during development, it is not surprising that disrupted ITP signaling causes developmental defects in *Tribolium* (Begum *et al*., 2009) and *Drosophila* (McEwan and Zandawala, unpublished). With regards to innate immunity, Gyc76C expression in the both the fat body and hemocytes is required for defense against Gram-positive bacteria (Iwashita *et al*., 2020). It remains to be seen if *ITP* knockdown also compromises immunity against Gram-positive bacteria. Besides previously identified functions of Gyc76C, our extensive expression mapping of this receptor also provides insights on other functions of ITPa-Gyc76C signaling. For instance, Gyc76C is expressed in female IPCs (Fig. 5 Supplement 2H), larval ring gland (Fig. 5 Supplement 1I) and the adult corpora allata (Fig. 5O). ITPa-Gyc76C signaling to the IPCs could modulate metabolic physiology associated with female reproduction. Moreover, ITPa-Gyc76C signaling could regulate juvenile hormone signaling in *Drosophila.* It could act similarly to its crustacean homolog, mandibular organ-inhibiting hormone (MOIH), which inhibits secretion of methyl farnesoate, a member of the juvenile hormone family, from the mandibular organs (Webster *et al*., 2012). This could in turn impact ovary development and/or vitellogenesis. ITP is also homologous to MIH, which inhibits ecdysteroid production by the Y-organs in crustaceans (Webster *et al*., 2012). Expression of Gyc76C in the larval ring gland, which also includes the ecdysteroid-producing prothoracic glands, suggests that ITP could regulate *Drosophila* development, as shown previously in *Tribolium* (Begum *et al*., 2009). It would also be of interest to determine the functions of Gyc76C in glia, especially those expressing the clock protein, Period (Fig. 5L). ITPa released by 5^th^-LN_v_ and LN_d_^ITP^ may link the neuronal clock with the clock in glial cells. Future studies could knockdown Gyc76C in these additional targets to identify novel roles of ITPa-Gyc76C signaling in *Drosophila*.

### Functional overlap between mammalian atrial natriuretic peptide (ANP) and Drosophila ITP

The multifunctional ITP signaling characterized here is reminiscent of the ANP signaling in mammals (Komatsu *et al*., 1991, Moro and Smith, 2009, Verboven *et al*., 2017). ANP is secreted from the cardiac muscle cells to regulate sodium and water excretion by the kidney. Interestingly, ITPL2 is also expressed in the heart muscles (Fig. 3M) and acts as an anti-diuretic in some contexts (Xu *et al*., 2023). Additional functions of ANP include roles in metabolism, heart function and immune system. Thus, ANP targets white adipocytes to affect lipid metabolism (Verboven *et al*., 2017) similar to the ITPa actions on the *Drosophila* fat body. Furthermore, ANP regulates glucose homeostasis, food intake and pancreatic insulin secretion as shown here for ITPa. A role of ANP as a cytokine in immunity, and with protective effects in tumor growth has also been implicated (De Vito, 2014), similar to the cytokine-like action of tumor-derived *Drosophila* ITPL2 (Xu *et al*., 2023). It is interesting to note that ITP and Gyc76C are absent in mammals and no orthologs of ANP have been discovered in invertebrates. While an ortholog of mammalian ANP receptors is present in *Drosophila*, studies characterizing its functions are lacking. It is possible that the ANP system in mammals acquired additional functions that are served by ITP signaling in invertebrates. Functional studies on *Drosophila* ANP-like receptors could shed light on the evolution of these signaling systems.

### Limitations of the study

It is worth pointing out that our phylogenetic analysis identified a second orphan mGC, Gyc32E, as a putative ITPa receptor. Although tissue expression analysis suggests that this receptor is not suited to mediate the osmoregulatory effects of ITPa, we cannot completely rule out the possibility that it also contributes to the metabolic phenotypes of ITPa via actions on IPCs and/or the fat body. It is also of interest to determine whether the three ITP splice forms act in synchrony in cases where they are colocalized in neurons. However, we were unable to determine the specific functions of ITPL1 and ITPL2, as existing RNAi transgenes from *Drosophila* stock centers target all three isoforms. Although ITPL2 functions as an anti-diuretic in a gut tumor model (Xu et al., 2023), the functions of ITPL2 released from the nervous system and under normal conditions are still unknown. Recent work in *Aedes aegypti* mosquitoes suggests that ITP and ITPL could have different functions (Sajadi and Paluzzi, 2024).

### Concluding remarks

To conclude, our comprehensive characterization of ITP, a homesotatic signaling system with pleiotropic roles, provides a foundation to understand the neuronal and endocrine regulation of thirst-driven behaviors and physiology. L-NSC^ITP^, with the potential to release up to eight diverse neuropeptides, likely regulate most aspects of *Drosophila* physiology to modulate systemic homeostasis.

## Acknowledgements

The authors would like to thank Irina Wenzel for helpful feedback during the preparation of this manuscript, Dr. Christian Wegener for use of the FlyPad facility, and Dr. Nils Reinhard for assistance with data analyses. We are also thankful to Francesca McEwan for preliminary analyses and to Manpreet Kooner, Selina Hilpert and Emilia Derksen for technical assistance. We thank the Princeton FlyWire team and members of the Murthy and Seung labs, as well as members of the Allen Institute for Brain Science, for development and maintenance of FlyWire (supported by BRAIN Initiative grants MH117815 and NS126935 to Murthy and Seung). M.Z. was supported by funding from the Deutsche Forschungsgemeinschaft (DFG; ZA1296/1-1), and NV INBRE grant from the National Institute of General Medical Sciences (GM103440). J.P.P was supported by a Natural Sciences and Engineering Research Council of Canada (NSERC) Discovery Grant and an Ontario Ministry of Research Innovation Early Researcher Award. A.B.B. and M.H.O. were supported by a NIGMS COBRE award P20GM130459. S.K. was supported by JSPS KAKENHI (20H03246). D.R.N was supported by funding from the Swedish Research Council (Grant Number: 2015-04626). F.S. was supported by NSERC CGS-D. J.G. was supported by funding from the University of Würzburg. We also acknowledge funding from the DFG for the Leica TCS SP8 microscope (251610680, INST 93/809-1 FUGG).

## Author contributions

D.R.N., J.P.P., and M.Z. conceived the study. J.G., D.R.N., M.O., J.P.P., and M.Z. supervised the project. J.G., F.S., M.A., H.N., A.B., E.D., S.H., S.K., L.T., M.O., J.P.P., and M.Z. performed the experimental work and analyzed the data. T.H.M. and M.Z. performed computational analyses. D.R.N., J.P.P and M.Z. wrote the manuscript. All authors read, provided feedback, and approved the final manuscript.

## Competing interest statement

We declare we have no competing interests.

## Data availability

Custom code used for connectome and single-cell transcriptome analyses is available at: https://github.com/Zandawala-lab/Gera-et-al-2025-Drosophila-ITPa-Gyc76C.

## Materials and Methods

### Fly strains

*Drosophila melanogaster* strains used in this study are listed in Supplementary Table 1. Unless stated otherwise, flies were raised at 25°C on a standard medium containing 8.0% malt extract, 8.0% corn flour, 2.2% sugar beet molasses, 1.8% yeast, 1.0% soy flour, 0.8% agar and 0.3% hydroxybenzoic acid. For adult-specific manipulations with *tubulin-GAL80[ts]*, flies were raised at 18°C until two days post-eclosion and then maintained at 29°C until analysis. Unless specified otherwise, all experiments were done using mated females.

### Immunohistochemistry and confocal imaging

Adult *Drosophila* were fixed in 4% paraformaldehyde (PFA) with 0.5% Triton-X100 in 0.1 M sodium phosphate buffer saline (PBST) for 2.5 hours on nutator at room temperature. Larval *Drosophila* were fixed in 4% PFA for 2 hours over ice. After fixation, the flies were washed with 0.5% PBST for 1 hour (4 x 15 mins). Subsequently, the flies were washed with PBS for 10 minutes. Fixed flies were then dissected in PBS and transferred to tubes containing blocking solution (5% normal goat serum in PBST with sodium azide at 1:100 dilution) on ice. After dissection, tissues were incubated in the primary antibody solution (diluted in blocking solution) for 48 hours at 4°C, followed by four washes with 0.5% PBST (4 x 15 mins), and incubated in secondary antibody (diluted in blocking solution) for 48 hours at 4°C. All the antibodies and fluorophores used in this study are listed in Supplementary Table 2. Finally, the flies were washed with 0.5% PBST (3 x 15 mins) followed by washes with PBS (2 × 10 mins). Samples were mounted using Fluoromount-G^TM^ (Invitrogen, Thermo Fisher) and imaged with a Leica SPE and TCS SP8 confocal microscopes (Leica Microsystems) using 20X glycerol, 40X oil or 63X glycerol immersion objectives.

### Fluorescence quantification

Confocal images were processed and the immunofluorescence levels measured using Fiji software. The final immunofluorescence of each sample was calculated by subtracting the background mean intensity from the mean intensity of the desired area.

### Sequence alignments and phylogenetic analysis

BLAST (Altschul *et al*., 1990) and HMMER (Potter *et al*., 2018) searches were performed using the *Drosophila* ITPa prepropeptide sequence to identify ITPL sequences in non-arthropods. ITP prepropeptide sequences were aligned using Clustal Omega (https://www.ebi.ac.uk/Tools/msa/clustalo/) and the conserved residues (at least 70% conservation) shaded using Boxshade (https://junli.netlify.app/apps/boxshade/). Phylogenetic analysis was performed using a custom workflow at NGPhylogeny.fr (Lemoine *et al*., 2019). Briefly, membrane guanylate cyclase receptor protein sequences (accession numbers for the sequences are included in the figure) were aligned using MAFFT (flavor: linsi; gap extension penalty: 0.123; gap opening penalty: 1.53; PAM 250 matrix). The alignment was trimmed using BMGE (BLOSUM 62 matrix; sliding window size: 3; maximum entropy threshold: 0; gap rate cut-off: 0.5; minimum block size: 5). A maximum-likelihood analysis with Smart Model Selection (model selection criteria: AIC; bootstrap: 500; random trees: 5) was used to generate the phylogeny. *Drosophila* guanylyl cyclase alpha and beta subunits were used as outgroups.

### Single-cell transcriptome analysis

Single-nucleus transcriptomes of fat body and Malpighian tubules were mined using the Fly Cell Atlas datasets (Li *et al*., 2022). Single-cell transcriptomes of *ITP*-expressing neurons were mined using the datasets generated earlier (Davie *et al*., 2018, Allen *et al*., 2020).

The parameters used to identify the different cell types are provided below:

LN^ITP^ (8 cells): ITP > 1 & NPF > 1 & cry > 0 & Phm > 0
L-NSC^ITP^ (7 cells): Tk > 1 & sNPF > 1 & ITP > 1 & ImpL2 > 1 & Crz == 0
L-NSC^DH31^ (6 cells): ITP > 2 & Dh31 > 4 & amon > 0 & Phm > 0
iag (1 cell): AstA > 0 & CCAP > 0 & ITP > 1 & Phm > 0 & amon > 0)
non-iag (23 cells): AstA == 0 & CCAP == 0 & ITP > 1 & Phm > 0 & amon > 0)
All analyses were performed in R-Studio (v2022.02.0) using the Seurat package (v4.1.1 (Hao *et al*., 2021)).

### Recombinant ITPa generation

ITP-PE (ITPa) was amplified from *w^1118^* adult mixed-sex whole body cDNA using forward (5’-gccaccATGTGTTCCCGCAACATAAAGATC-3’) and reverse (5’-GCACTTTACTTGCGACCCAGG-3’) gene-specific primers and cloned into pGEM T-easy vector and sub-cloned into pcDNA3.1+ mammalian expression vector using standard molecular techniques as previously described (Wahedi and Paluzzi, 2018). Recombinant ITPa was expressed in AtT-20 cells (ATCC CCL-89), which is a murine-derived cell line of neuroendocrine origin from pituitary tumour, by transfection using Lipofectamine LTX reagent following the manufacturer’s protocol. A pcDNA3.1+ vector containing mCherry instead of the ITPa construct was used as a control to monitor transfection efficiency. A stable cell line constitutively expressing ITPa was isolated under selection using 600µg/mL geneticin and scaled up to yield recombinant ITPa for *ex vivo* Ramsay assay. Heterologous expression of ITPa was verified by immunoblot using a rabbit polyclonal antiserum against the C-terminal region of *Drosophila* ITPa described previously (Hermann-Luibl *et al*., 2014, Galikova *et al*., 2018) diluted 1:8000 in immunoblot block buffer, whereas E7 beta-tubulin (1:2500) was used as loading control (deposited to the DSHB by Klymkowsky, M.; DSHB Hybridoma Product E7) following a previously described immunoblot protocol (Rocco and Paluzzi, 2020). This confirmed ITPa expression in AtT-20 cells while no such band was detected in mCherry expressing cells (Fig. 6 Supplement 2). Cell lysates were collected and protein samples semi-purified by size-exclusion filtration using centrifugal concentrators with a polyethersulfone membrane (ThermoFisher Scientific, Waltham, MA). Specifically, protein harvested from AtT-20 cells expressing ITPa was centrifuged through 20 kDa molecular weight cut off (MWCO) concentrators and the flow through excluding proteins >20 kDa was then transferred to a second centrifugal concentrator with a 5 kDa MWCO. This allowed the expressed ITPa to be concentrated in the retentate since its molecular weight is ∼9 kDa and permitted buffer exchange so that the final semi-purified ITPa was reconstituted in 1x phosphate buffered saline (PBS). The concentration of the semi-purified ITPa was determined by an indirect enzyme-linked immunosorbent assay as previously described (MacMillan *et al*., 2018) using the C-terminal antigen used to generate the ITPa antiserum as a standard.

To improve purity of heterologously expressed ITPa and to scale up production, recombinant ITPa was independently produced by Genscript (Genscript, Piscataway, NJ) following heterologous expression in proprietary TurboCHO™ and TurboCHO™ 2.0 expression systems (Genscript, Piscataway, NJ). To produce C-terminally amidated recombinant ITPa (ITP-PE), human peptidylglycine alpha-amidating monooxygenase was co-expressed along with ITP-PE in the expression system. ITPa included an N-terminal histidine tag that allowed one-step purification following heterologous expression.

### *Ex vivo* fluid secretion (Ramsay) assay

Fluid secreted by individual Malpighian tubules was monitored using the classical Ramsay assay (Ramsay, 1954) where adult fly Malpighian tubule secretion rates were measured following protocols recently described in detail (MacMillan *et al*., 2018). Briefly, adult male flies (5-6 days old) were dissected under *Drosophila* saline (Vanderveken and O’Donnell, 2014) and the anterior pair of Malpighian tubules was isolated from the gut at the ureter and then transferred into a 20 µl droplet (comprised of a 1:1 mixture of Schneider’s insect medium and *Drosophila* saline) placed over a small well within a Sylgard-lined Petri dish filled with hydrated paraffin oil to prevent sample evaporation. The proximal end of a single Malpighian tubule was pulled out of the bathing droplet and wrapped around a minuten pin so that the ureter was approximately halfway between droplet and the pin. As the Malpighian tubule incubates in the bathing droplet, a secretory droplet forms at the ureter, which following a 60 min incubation, is then detached and measured using a calibrated eyepiece micrometer. The volume of the secreted fluid is then calculated using the secreted droplet’s diameter that allows the fluid secretion rate (FSR) to be determined (FSR = droplet volume/incubation time). To stimulate fluid secretion, diuretic hormones including *Drosophila* leucokinin and DH_31_ were added into the bathing droplet to achieve a final concentration of 10nM and 1μM, respectively. Unstimulated tubules were treated with a 1:1 mixture of Schneider’s insect medium and *Drosophila* saline alone or with diluted PBS for experiments involving recombinant ITPa.

### GPCR heterologous assay

*Drosophila* GPCRs PK2R (CG8784) and DTKR (CG7887), which are homologous to *B. mori* ITPa and ITPL receptors, respectively (Nagai *et al*., 2014), were amplified using gene-specific primers as previously described (Park *et al*., 2002, Birse *et al*., 2006) and sub-cloned into the pcDNA3.1^+^ using standard molecular biology techniques. Receptors were expressed in CHO-K1 cells stably expressing aequorin (CHOK1-aeq), a calcium-activated bioluminescent protein (Sajadi *et al*., 2020). At 48 hrs post transfection with either PK2R or DTKR, CHOK1-aeq cells were prepared for the heterologous functional assay by resuspension in BSA assay media (DMEM-F12 media containing 0.1% bovine serum albumin (BSA), 1X antimycotic-antibiotic) containing 5µM coelenterazine *h* (Nanolight Technologies, Pinetop, AZ, USA) and incubated with mixing for three hours. After this incubation, cells were diluted 10-fold with BSA assay media reducing the concentration of coelenterazine *h* to 0.5µM and incubating for an additional hour with constant mixing. Cells were then loaded into individual wells of a white 96-well luminescence plate with an automatic injector unit and luminescence was measured for 20 seconds using a Synergy 2 Multi-Mode Microplate Reader (BioTek, Winooski, VT, USA). Each well of the 96-well plate was pre-loaded with candidate ligands (recombinant ITPa, pyrokinin 2 and tachykinin 1) at 100nM and 500nM (Park *et al*., 2002, Birse *et al*., 2006). BSA assay media alone was utilized as a negative control while 50µM ATP, which acts on endogenously expressed purinoceptors (Iredale and Hill, 1993), was used as a positive control.

### Gyc76C characterization in HEK293T cells

*Drosophila* Gyc76C, codon-optimized for mammalian expression, was custom-synthesized and sub-cloned into pCDNA3.1(+)-C-HA by Genscript (Piscataway, NJ). HEK293T cells were cultured in Dulbecco’s modified Eagle’s medium (Corning) supplemented with 10% fetal bovine serum (GenClone, El Cajon, CA) at 37°C and 5% CO_2_. Cells plated in 35mm glass bottom imaging dishes (Cellvis, Mountain View, CA) were transfected at 70-80% confluency with 1.5-1.75 µg Gyc76C_pCDNA3.1(+)-C-HA and/or 0.75-1.0 µg Green cGull (Matsuda *et al*., 2017) using Mirus TransIT-LT1 (Mirus Bio, Madison, WI) following manufacturer’s protocol. Cells were then incubated at 30.5°C and 5% CO_2_ for 24 hours. For imaging, HEK293T media was replaced with modified Ringer’s buffer (140mM NaCl, 3.5mM KCl, 0.5mM NaH_2_PO_4_, 0.5mM S-3 MgSO_4_, 1.5mM CaCl_2_, 10mM HEPES, 2mM NaHCO_3_, and 5mM glucose) 30 minutes before imaging. Live-cell images were acquired every 15 s for 6 min using a Stellaris X8 confocal microscope with an 86X water objective. After 90 s of baseline recording, recombinant ITP-PE/PAM (Genscript) was added at 50nM, 250nM, or 500nM final concentration.

Image analysis was performed using FIJI (ImageJ) software. Regions of interest (ROIs) were drawn around cells expressing Green cGull and fluorescence intensity was measured at every timepoint through the stack. The same ROIs were then moved to off-cell regions of the image and background measurements were taken at every timepoint. Background was subtracted from cell measurements in Excel. Baseline was set to the average of the first 7 timepoints and all subtracted measurements were divided by this value. Replicates were compiled in GraphPad Prism 10 software for visualization. Area under the curve was computed as the sum of fluorescence intensity over baseline for each cell and then compiled using Prism 10 for visualization. Statistical analysis was performed using nonparametric distribution one-way ANOVA without matching. Post-hoc multiple comparisons were corrected and subjected to Dunn’s test.

To assess Gyc76C expression in imaged cells, imaging dishes were fixed in 4% PFA and subsequently stained with HA-Tag Rabbit mAb (1:2000; Cell Signaling Technology, Danvers, MA) and anti-GFP Goat pAb (1:1000; Rockland). Secondary antibodies (all at 1:1000) included donkey-anti-goat highly cross-absorbed IgG Alexa Fluor 488 (Invitrogen, Carlsbad, CA) or donkey anti-goat IgG Star Green (Abberior, Göttingen, DE) and donkey anti-rabbit highly cross-absorbed IgG Alexa Fluor 647 (Invitrogen, Carlsbad, CA). Fixed cells were imaged on a Keyence BzX-710 microscope with a 20X objective. Images were processed using FIJI (ImageJ) software. Correlations between stained signal intensity and peak live Green cGull fluorescence were performed using Prism 10.

### Feeding assays

flyPAD (Itskov *et al*., 2014) was used to calculate the number of feeding bouts over 24 hours as well as preference between sucrose versus yeast. Individual flies were mouth-pipetted to a flyPAD unit and were given a choice between sucrose (5mM) and yeast (10%) in 2% agarose. The data was analyzed using a custom script provided by the manufacturer. Total food intake over 24 hours was calculated by adding the number of feeding bouts on sucrose and yeast. Food preference for each fly was calculated by dividing the difference in sucrose and yeast uptake with total food intake.

CAFE assay was performed to monitor the preference between nutritive and non-nutritive sugars. For each genotype, 10 flies (fed or starved) were transferred to an empty glass vial and given a choice between 25mM D-fructose (nutritive) and 80mM D-arabinose (non-nutritive) using 5μL capillaries. Glass vials containing food capillaries without flies were used as controls to monitor evaporation. All the glass vials were maintained in moist chambers and reduction in the volume of individual capillary was measured after 2 hours of feeding. Food preference was calculated as above based on 20 replicates for each genotype.

For experiments involving *ITP* knockdown, total feeding was analysed using the CAFE assay. Ad-libitum fed female flies were anesthesized with CO_2_ and individually transferred into 2ml tubes. Each fly was provided a 5 µl glass capillary filled with a solution containing 10 % sucrose, 10 % yeast and 0.1 % propionic acid. Flies were allowed to feed for 24 hours before measuring the volume they consumed. To reduce evaporation, all tubes were kept in a humidified box at 25°C on a 12:12 light dark cycle.

### Glucose assay

Thorax of around 40-50 adult flies were punctured using a 0.1mm metallic needle. The flies were then transferred to 0.5ml tubes with a hole at the bottom. These tubes containing the flies were placed inside a 1.5ml tube and centrifuged for 10 minutes at 5000 RPM at 4°C. The clear hemolymph collected in the 1.5ml tubes was used to measure glucose concentration as per the manufacturer recommended protocol (Glucose calorimetric assay kit, Cayman #10009582). A minimum of 10 replicates were analyzed for each genotype.

### Triglyceride assay

To quantify total triglycerides, 5 flies for each genotype were homogenized and processed as per the manufacturer’s protocol (Triglyceride Calorimetric assay kit, Cayman #10010303). Triglyceride levels were normalized by the protein content. A minimum of 9 replicates were analyzed for each genotype.

### Glycogen assay

To quantify the amount of stored glycogen, 5 flies for each genotype were homogenized and processed as per the manufacturer’s protocol (Glycogen assay kit, Cayman #700480). The amount of glycogen was normalized by the protein content. A minimum of 10 replicates were analyzed for each genotype.

### Protein content

Protein concentrations were measured using the Bradford reagent (Sigma #B6916). Samples were added at a 1:200 ratio in a 96-well plate and incubated at room temperature for 5 minutes. Finally, the absorbance was measured at 595nm using a Magellan Sunrise plate reader.

### Water content

Groups of five flies were anesthetized with CO_2_, placed into tubes and frozen at −80°C for a few hours. Wet weight of the flies was first determined by weighing them. The flies were then dried at 65°C for 48 hours before determining their dry weight. The water content was calculated by subtracting dry weight from wet weight.

### Defecation assay

Flies were anesthetized with CO_2_ and individually placed into small glass tubes containing blue food (4% sucrose, 2% agarose and 1% blue food coloring (“Brilliant blue FCF”)) for two overnights at 25°C. Afterwards, individual flies were flipped into empty glass tubes, which were placed horizontally in a box at 25°C. The number of feces droplets in each glass tube were manually counted at 2-hour intervals for up to six hours.

### *Drosophila* activity monitoring (DAM) experiments

To monitor the locomotor activity of individual flies, *Drosophila* activity monitoring system (Trikinetics Inc., Waltham, Massachusetts) was used. Individual flies were transferred to a thin glass tube (length 5cm, diameter 5mm) containing 2% agar and 4% sucrose for fed conditions, 2% agar for starved conditions or left empty for desiccating conditions. Activity was recorded in 1-minute intervals under 12:12 light dark cycles for 8-10 days followed by 8-10 days of constant darkness. The light dark cycles were maintained using the LED light sources set at 100 lux, housed in a chamber maintained at constant temperature and 70% relative humidity ± 5%. The data was analyzed using Actogram J in Fiji (Schmid *et al*., 2011). All analyses were based on approximately 30 flies per genotype.

To assess the impact of desiccation on locomotor activity, locomotor activity of individual flies was recorded in tubes containing 2% agar and 4% sucrose for approximately 66 hours. Following this time, flies were transferred to empty tubes to assess the impact of desiccation on locomotor activity.

### Stress tolerance assays

To monitor starvation survival, flies were individually placed in glass tubes containing 2% agar and their survival estimated automatically (based on lack of activity) using the DAM system as above. To monitor survival under desiccation, groups of 20 flies were kept in empty vials without access to any water or food. Dead flies were quantified visually at regular intervals during daytime. Survival curves were generated based on at least 120 flies per genotypes. Tolerance to salt stress was monitored by maintaining groups of 20 flies each on an artificial diet (medium containing 100 g/L sucrose, 50 g/L yeast, 12 g/L agar, 3 ml/L propionic acid and 3 g/L nipagin) supplemented with 4% NaCl. Number of dead flies were quantified visually at regular intervals during daytime. Survival curves were generated based on at least 120 flies per genotypes. To assess recovery from chill coma, 10 flies for each genotype were transferred into empty vials and kept in ice cold water (0°C) for 4 hours to induce immediate chill coma. Following this incubation, the vials were transferred to room temperature and the recovery of flies monitored visually at 2 min intervals. Approximately 100 flies per genotype were analyzed.

### Ovary imaging

Around 50-60 ovaries of each genotype were fixed, mounted, and imaged using a bright field microscope.

### Synaptic connectivity analyses and data visualization

ITPa-expressing neurons in the FlyWire brain connectome were identified previously (McKim *et al*., 2024, Reinhard *et al*., 2024). FlyWire cell IDs of identified ITPa neurons are provided in Supplementary Table 3. We used the v783 snapshot of the FlyWire connectome and its annotations for all the analyses (Dorkenwald *et al*., 2024, Schlegel *et al*., 2024). Connectivity was based on updated synapse predictions (Yu *et al*., 2025). We used a threshold of 5 synapses to identify significant connections. Connectivity analyses were based on custom scripts generated previously (McKim *et al*., 2024, Reinhard *et al*., 2024). All data for figure visualizations were processed and analyzed in R-Studio (2024.04.2+764). FlyWire neuroglancer was used to visualize neuron reconstructions (Dorkenwald *et al*., 2022).

### Statistical analyses

Unless mentioned otherwise, an unpaired t-test was used for comparisons between two genotypes and one-way analysis of variance (ANOVA) followed by Tukey’s multiple comparisons test for comparisons between three genotypes. The horizontal line in box-and-whisker plots represents the median. Log-rank (Mantel-Cox) test was used to compare survival and chill coma recovery curves. All statistical analyses were performed using GraphPad Prism and the confidence intervals are included in the figure captions.

## Notes

### Competing Interest Statement

The authors have declared no competing interest.

### Summary of Updates

This version of the manuscript includes substantial revisions based on feedback from the peer review process at eLife. The revisions include new figures based on new experiments, restructuring of original text and figures to accommodate new results, expanded discussion, additional methods and additional coauthors. To strengthen our findings on ITPa signaling via Gyc76C, this version includes experimental data showing Gyc76C activation by ITPa in a heterologous system. It also includes a figure showing phenotypes following knockdown of ITP in ITPa-producing cells. Additional data showing the effects of Gyc76C knockdown using an independent RNAi construct are included in the supplementary material. Calcium activity of ITPa-expressing neurons (measured using the CaLexA system) under different contexts is now included as a new figure. The connectomic analyses have been updated using an improved connectome dataset that was released during the revision of this manuscript. A broader discussion of Gyc76C and its known functions in Drosophila has now been included.

