## Supplementary material for "Anti-diuretic hormone ITP signals via a guanylate cyclase receptor to modulate systemic homeostasis in *Drosophila*"

**Supplementary Table 1:** Fly strains used in this study

| **Fly strain** | **Stock number / Reference** |
|---|---|
| *ITP-RC-GAL4* | (Deng *et al.*, 2019) |
| *ITP-RD-GAL4* | BDSC# 84702 |
| *yolk-GAL4* | BDSC# 58814 |
| *Uro-GAL4* | (Halberg *et al.*, 2016) |
| *Uro-GAL4; UAS-dicer* | Dr. Kenneth Halberg |
| *c724-GAL4* | (Feingold *et al.*, 2019) |
| *CCAP-GAL4* | BDSC# 25685 |
| *Gyc76C-GAL4* | (Kondo *et al.*, 2020) |
| *Gyc32E-GAL4* | BDSC# 81160 |
| *Lkr-GAL4* | (Zandawala *et al.*, 2018) |
| *PK2-R1-GAL4* | BDSC# 84686 |
| *CG30340-GAL4* | BDSC# 84611 |
| *TkR99D-GAL4* | BDSC# 76208 |
| *UAS-Gyc76C-RNAi #1* | VDRC# 106525 |
| *UAS-Gyc76C-RNAi #2* | BDSC# 57315 |
| *UAS-ITP-RNAi* | VDRC# 330029 |
| *UAS-ITPa* | (Hermann-Luibl *et al.*, 2014) |
| *UAS-myr::tdTomato;2xLexAop-GFP;UAS-CaLexA,LexAopGFP/TM6B, Tb (UAS-CaLexA)* | Dr. Kenneth Halberg |
| *JFRC81-10xUAS-IVS-Syn21-GFP-p10 (UAS-JFRC81GFP)* | (Pfeiffer *et al.*, 2012) |
| *JFRC29-10xUAS-IVS-myr::GFP-p10 (UAS-JFRC29GFP)* | (Pfeiffer *et al.*, 2012) |
| *UAS-nls-mCherry* | BDSC# 38425 |
| *Tubulin-GAL80[ts]* | BDSC# 7017 |
| $w^{1118}$ | BDSC# 5905 |
| *UAS-Luciferase-RNAi* | BDSC# 31603 |
| *VDRC RNAi control* | VDRC# 60000 |

**Supplementary Table 2:** Antibodies used for immunohistochemistry

| Antibody | Dilution | Source / Reference |
|---|---|---|
| ***Primary antibodies*** | | |
| chicken anti-GFP | 1:1000 | Abcam, RRID: AB_300798 |
| mouse anti-GFP | 1:1000 | Thermo Fisher Scientific |
| goat anti-GFP | 1:1000 | Rockland immunochemicals |
| guinea pig anti-GFP | 1:1500 | Synaptic systems (#132005) |
| rat anti-mCherry (16D7) | 1:1000 | Thermo Fisher Scientific |
| rabbit anti-ITPa | 1:5000 | Dr. Heinrich Dircksen (Hermann-Luibl *et al.*, 2014) |
| guinea pig anti-ITPa | 1:5000 | Dr. Heinrich Dircksen |
| rabbit anti-$DH_{31}$ | 1:1000 | Dr. Jan Veenstra (Park *et al.*, 2008) |
| anti-*Diploptera punctata* Ast-A7 | 1:2000 | Dr. Christian Wegener |
| rabbit anti-NPF (to label Tv neurons) | 1:1000 | Dr. Mark Brown |
| rabbit anti-AKH | 1:1000 | Dr. Mark Brown |
| rabbit anti-DILP2 | 1:2000 | Dr. Jan Veenstra (Veenstra *et al.*, 2008) |
| rabbit anti-PER | 1:1000 | Dr. Charlotte Helfrich-Förster (Stanewsky *et al.*, 1997) |
| mouse nc82 anti-Bruchpilot | 1:50 | Dr. Charlotte Helfrich-Förster (Wagh *et al.*, 2006) |
| rabbit anti-HA | 1:2000 | Cell Signaling Technology |
| ***Secondary antibodies and fluorescent stains*** | | |
| DAPI | 1:1000 | |
| Hoechst (20mM) | 1:1000 | |
| Nile red | 1.1000 | Sigma Aldrich |
| Rhodamine-phalloidin | 1:1000 | Thermo Fisher Scientific |
| donkey anti-guinea pig Alexa Fluor ® 488 | 1:1000 | Thermo Fisher Scientific |
| goat anti-chicken Alexa Fluor® 488 | 1:1000 | Thermo Fisher Scientific |
| goat anti-mouse Alexa Fluor® 488 | 1:1000 | Thermo Fisher Scientific |
| donkey anti-goat Alexa Fluor® 488 | 1:1000 | Thermo Fisher Scientific |
| donkey anti-goat Star Green | 1:1000 | Abberior |
| donkey anti-guinea pig Alexa Fluor ® 555 | 1:1000 | Thermo Fisher Scientific |
| donkey anti-rabbit Alexa Fluor ® 555 | 1:1000 | Thermo Fisher Scientific |
| goat anti-mouse Alexa Fluor® 555 | 1:1000 | Thermo Fisher Scientific |
| goat anti-guinea pig Alexa Fluor® 647 | 1:1000 | Thermo Fisher Scientific |
| donkey anti-rabbit Alexa Fluor ® 647 | 1:1000 | Thermo Fisher Scientific |
| donkey anti-mouse Alexa Fluor® 647 | 1:1000 | Thermo Fisher Scientific |
| donkey anti-rat Alexa Fluor® 555 | 1:1000 | Thermo Fisher Scientific |

**Supplementary Table 3:** Root IDs (v783) of ITPa-producing cells in the FlyWire connectome

| Neuron type | Root ID | Hemisphere |
|---|---|---|
| L-NSC$^{ITP}$ | 720575940630046506 | right |
| L-NSC$^{ITP}$ | 720575940629808911 | right |
| L-NSC$^{ITP}$ | 720575940629872971 | right |
| L-NSC$^{ITP}$ | 720575940627436035 | right |
| L-NSC$^{ITP}$ | 720575940625721118 | left |
| L-NSC$^{ITP}$ | 720575940629924091 | left |
| L-NSC$^{ITP}$ | 720575940631592017 | left |
| L-NSC$^{ITP}$ | 720575940613850262 | left |
| 5$^{th}$-LN$_v$ | 720575940619074049 | right |
| 5$^{th}$-LN$_v$ | 720575940625254636 | left |
| LN$_d$$^{ITP}$ | 720575940627933336 | right |
| LN$_d$$^{ITP}$ | 720575940634984800 | left |

A

| ***B.mori* receptor** | ***B. mori* ligand** | ***Drosophila* receptor** | ***Drosophila* ligand** |
|---|---|---|---|
| BNGR-A2 | ITPa | CG8784 (PK2-R1) | Pyrokinin |
| BNGR-A34 | ITPa | CG30340 | Unknown |
| BNGR-A24 | Tachykinin, ITPL | CG7887 (TkR99D) | Tachykinin, ITPL2 |

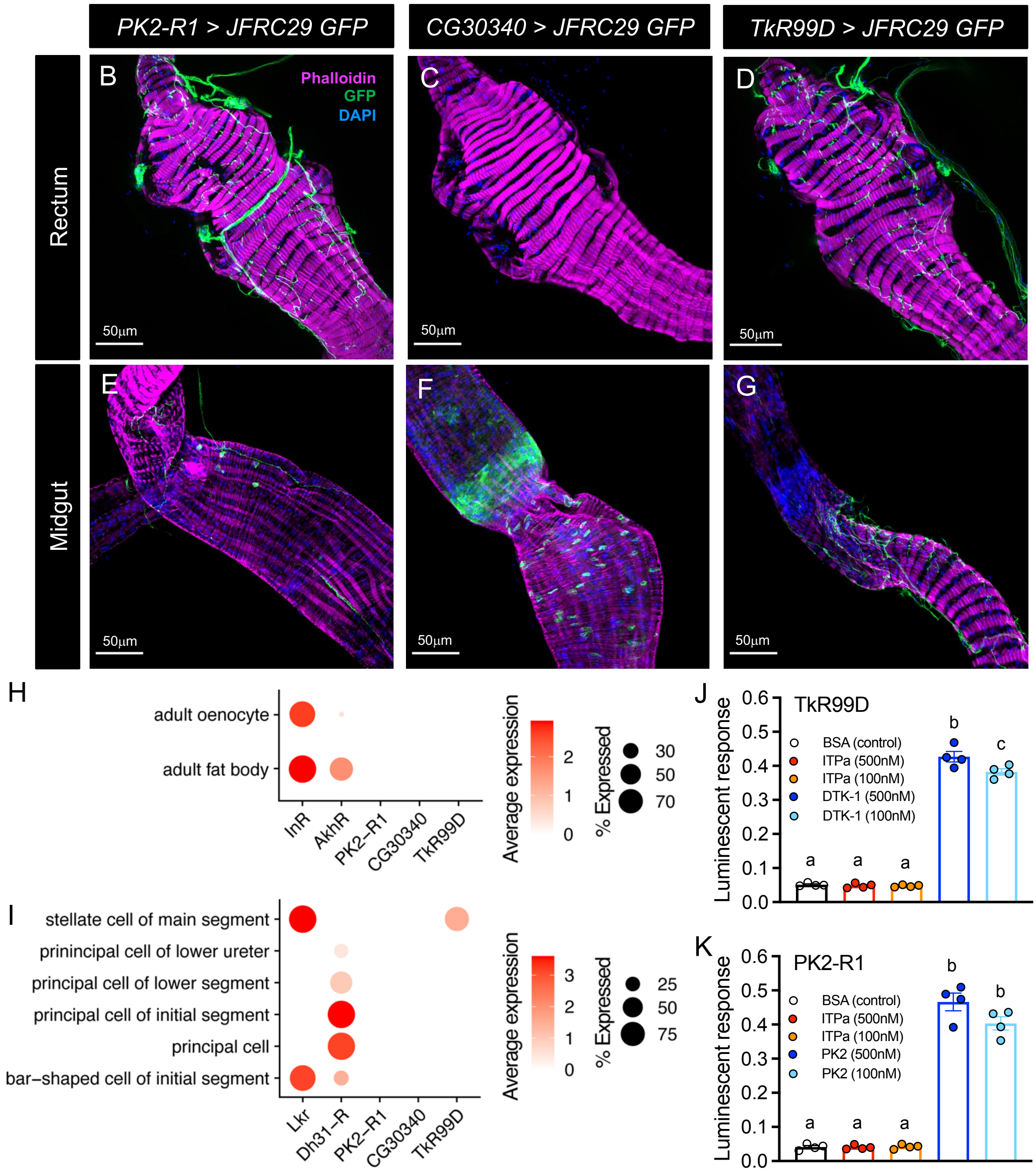


**Figure 4 Supplement 1: *Drosophila* orthologs of *Bombyx mori* ITP and ITPL receptors. (A)** *Drosophila* CG8784 (PK2-R1), CG30340 and CG7887 (TkR99D) are orthologs of *Bombyx mori* BNGR-A2, BNGR-A34 and BNGR-A24, respectively. Ligands of these receptors based on previous studies are indicated (Nagai *et al.*, 2014, Nagai-Okatani *et al.*, 2016). **(B)** PK2-R1 is not expressed in the rectal papillae but is expressed in the neurons innervating the rectum. **(C)** CG30340 is not expressed in the rectum. **(D)** Tkr99D is also only expressed in axons innervating the rectum. **(E)** PK2-R1, **(F)** CG30340 and **(G)** TkR99D are all expressed in the midgut or the neurons innervating it. **(H)** PK2-R1, CG30340 and TkR99D are not expressed in the fat body but receptors for insulin (InR) and adipokinetic hormone (AkhR) are. **(I)** PK2-R1 and CG30340 are not expressed in Malpighian tubules. TkR99D and leuckonin receptor (Lkr) are expressed in stellate cells and diuretic hormone 31 receptor (Dh31-R) is expressed in principal cells. **(J)** *Drosophila* tachykinin 1 (DTK-1) but not ITPa activates TkR99D expressed in CHO-K1 cells stably expressing aequorin (CHOK1-aeq), a calcium-activated bioluminescent protein. **(K)** Pyrokinin 2 (PK2) but not ITPa activates PK2-R1 expressed in CHOK1-aeq cells. For **J** and **K**, bars labeled with different letters are significantly different from each other ($p < 0.05$, as assessed by one-way ANOVA followed by Tukey's multiple comparisons test).

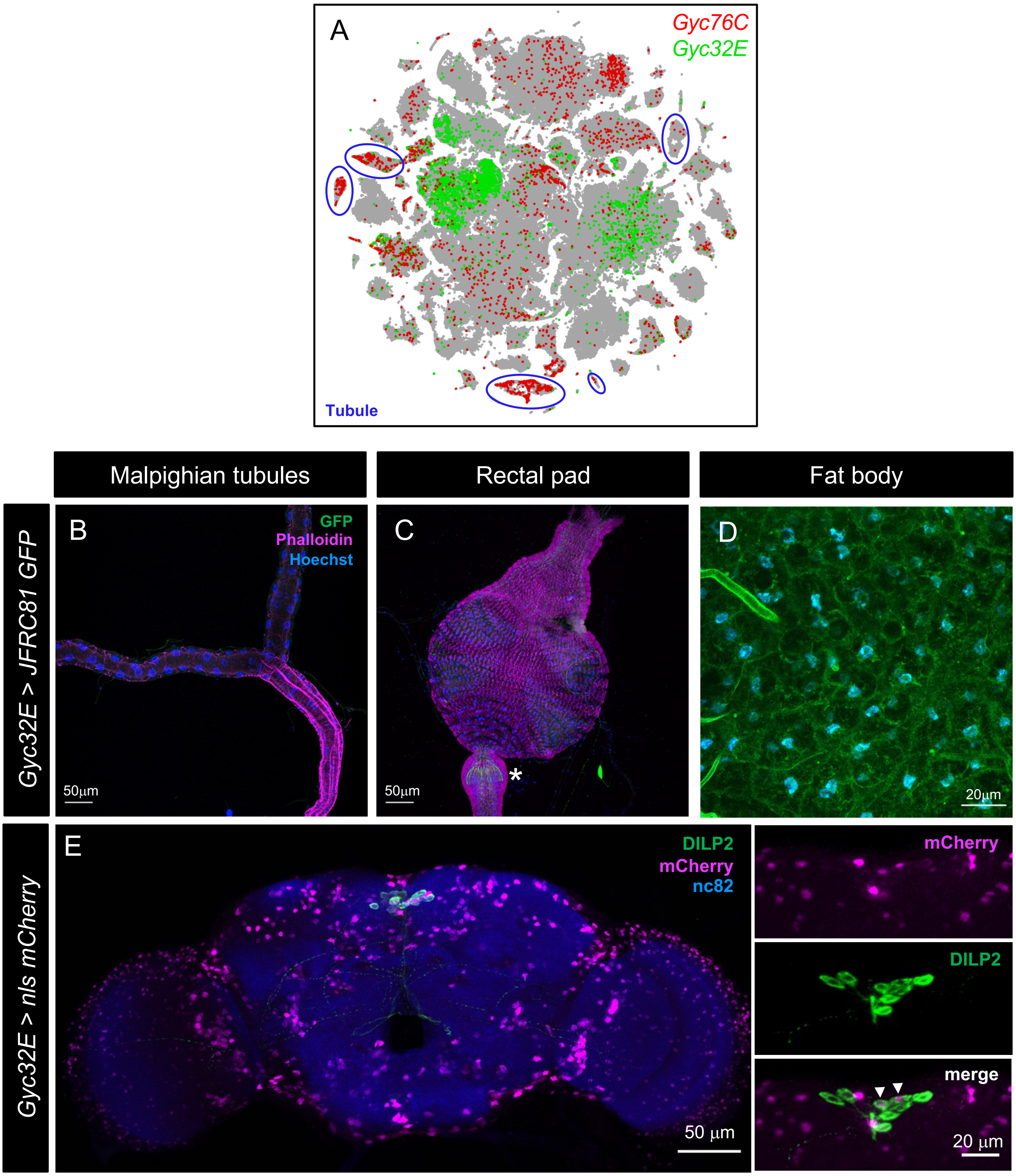


**Figure 4 Supplement 2: Expression of candidate ITPa/ITPL receptors in adult male *Drosophila* tissues. (A)** *Gyc76C* expression in Malpighian tubules (blue outline) is higher compared to *Gyc32E*. *Gyc32E-GAL4* does not drive GFP expression in the **(B)** Malpighian tubules and **(C)** rectal pad but is expressed in the hindgut (marked by an asterisk) and **(D)** fat body of males. **(E)** *Gyc32E-GAL4* drives nuclear mCherry expression in a subset of insulin-producing cells (labelled by DILP2 antibody).

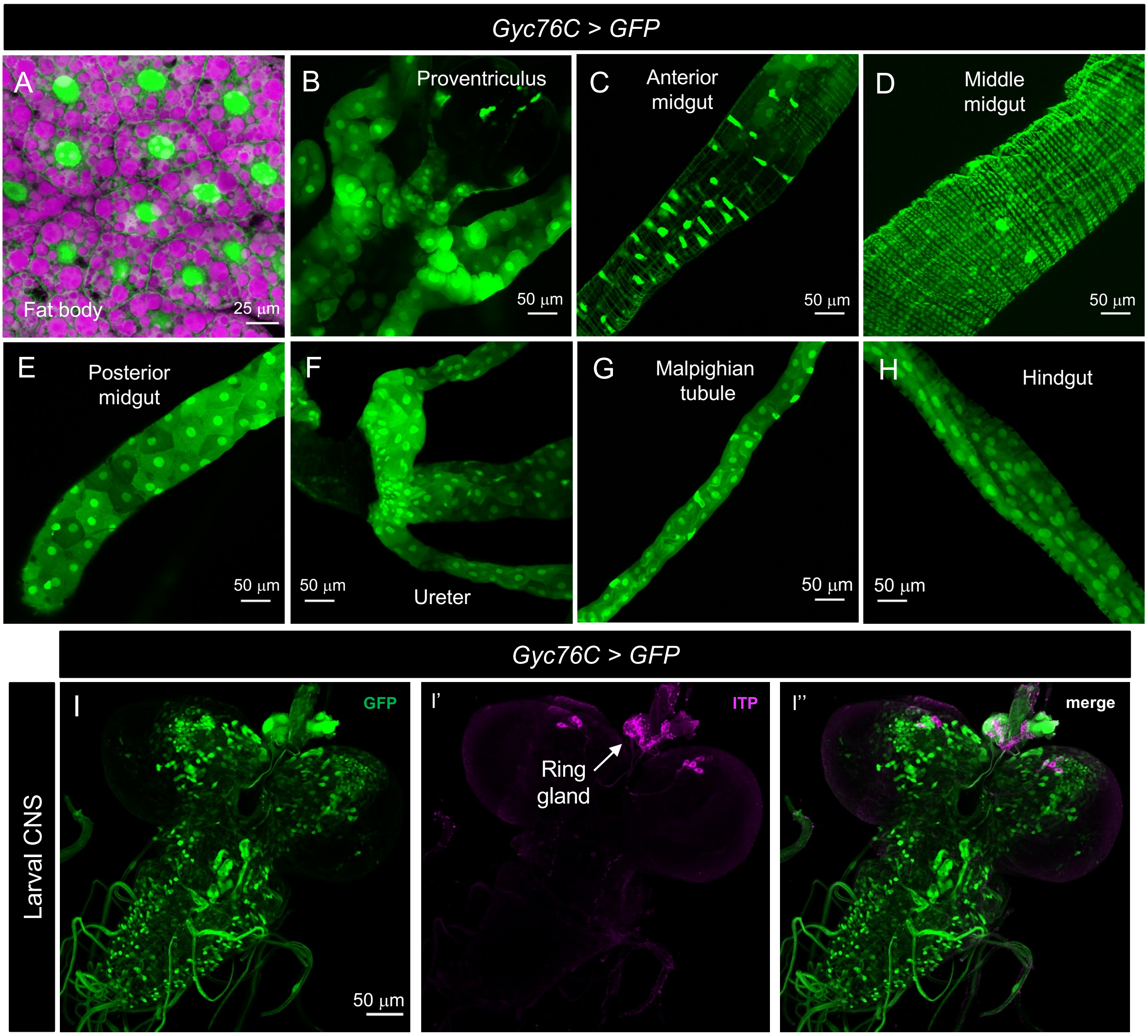


**Figure 5 Supplement 1: Gyc76C expression in larval *Drosophila*.** *Gyc76C-T2A-GAL4* drives GFP (UAS-JFRC81GFP) expression in the **(A)** fat body, **(B)** proventriculus, **(C)** anterior midgut, **(D)** middle midgut, **(E)** posterior midgut, **(F)** ureter, **(G)** Malpighian tubules and **(H)** hindgut. Lipid droplets (magenta in panel **A**) were stained using Nile red. **(I)** Gyc76C is broadly expressed in the larval nervous system, including the ring gland, which is innervated by ITPa-expressing neurons (magenta).

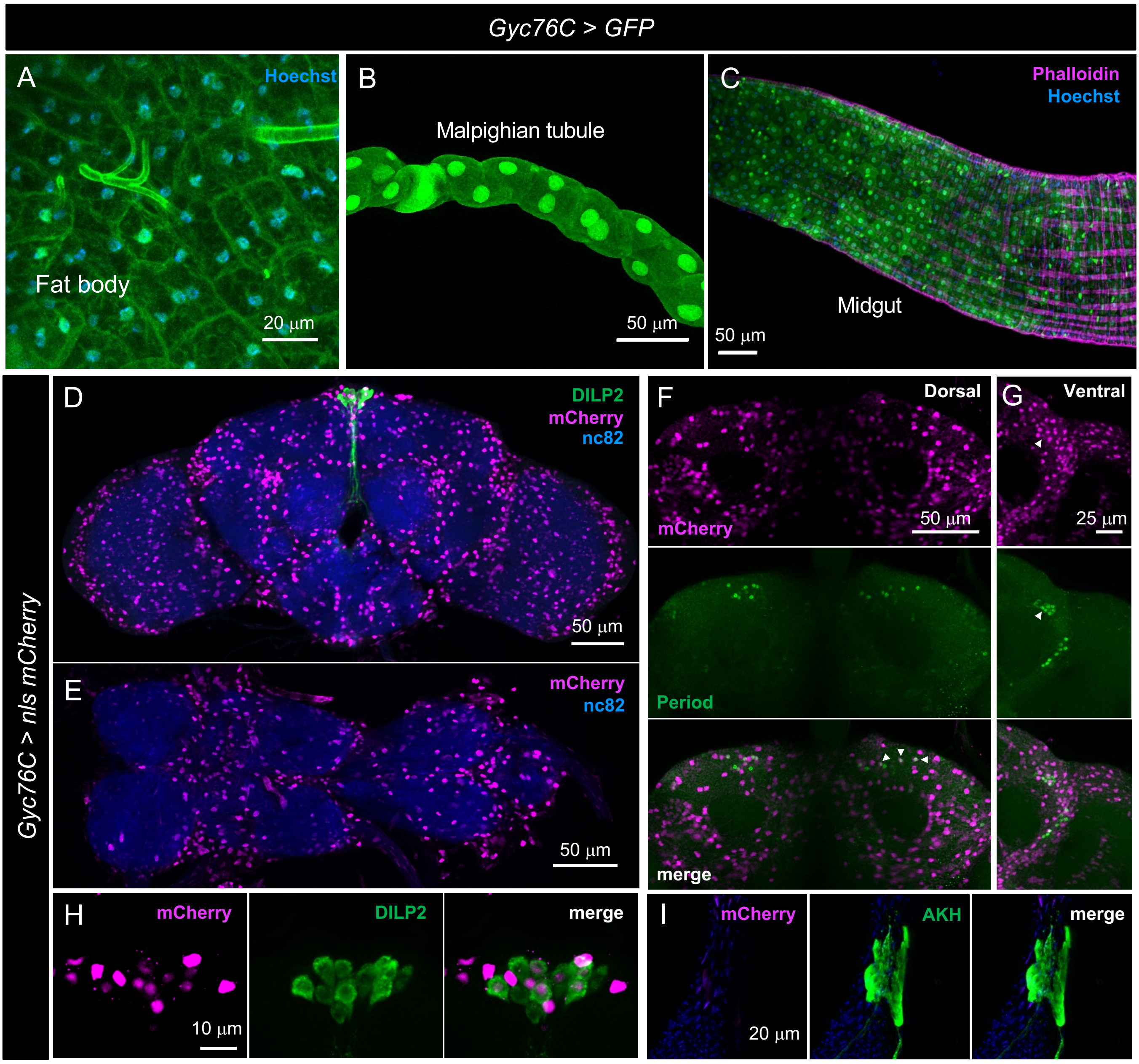


**Figure 5 Supplement 2: Gyc76c expression in adult female *Drosophila*.** *Gyc76C-T2A-GAL4* drives GFP (UAS-JFRC81GFP) expression in the **(A)** fat body, **(B)** Malpighian tubules and **(C)** the midgut. *Gyc76C-T2A-GAL4* drives nuclear mCherry expression in the brain **(D)** and ventral nerve cord **(E)**. **(F and G)** Gyc76C is expressed in subsets of clock neurons (labelled by Period antibody and marked by arrowheads). Gyc76C is expressed in **(H)** a subset of insulin-producing cells (labelled by DILP2 antibody) but not **(I)** in the adipokinetic hormone (AKH) producing endocrine cells.

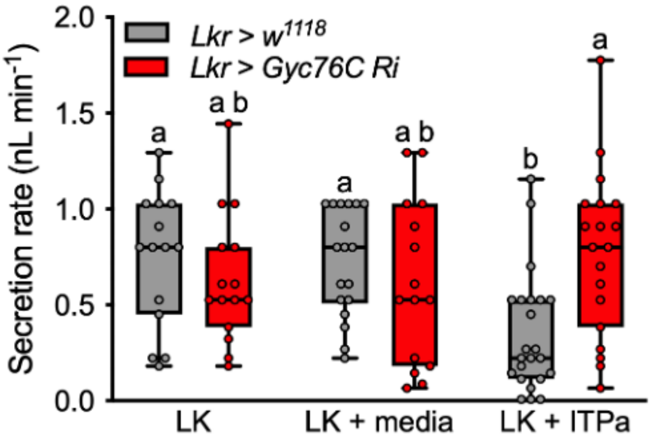


**Figure 6 Supplement 1: Recombinant *Drosophila* ITPa inhibits Malpighian tubule secretion via Gyc76C.** Recombinant ITPa expressed in AtT-20 cells inhibits leucokinin (LK)-stimulated secretion by renal tubules. This inhibitory effect is abolished in tubules where *Gyc76C* has been knocked down with *UAS-Gyc76C RNAi* (*#106525*) using the *LK receptor GAL4* (*Lkr-GAL4*). Differences between treatments within and across genotypes are denoted by different letters as determined by two-way ANOVA followed by Tukey's multiple comparisons test ($p < 0.05$).

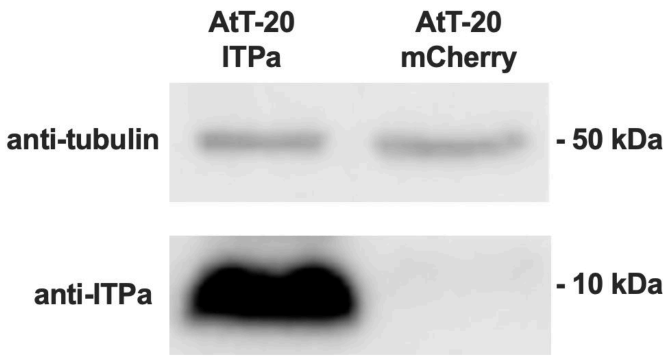


**Figure 6 Supplement 2: Western blot analysis of recombinant ITPa produced in AtT-20 cells.** A single band corresponding to the molecular weight of ITPa (~9 kDa) is observed in lysates from AtT-20 cells expressing ITPa but not mCherry.

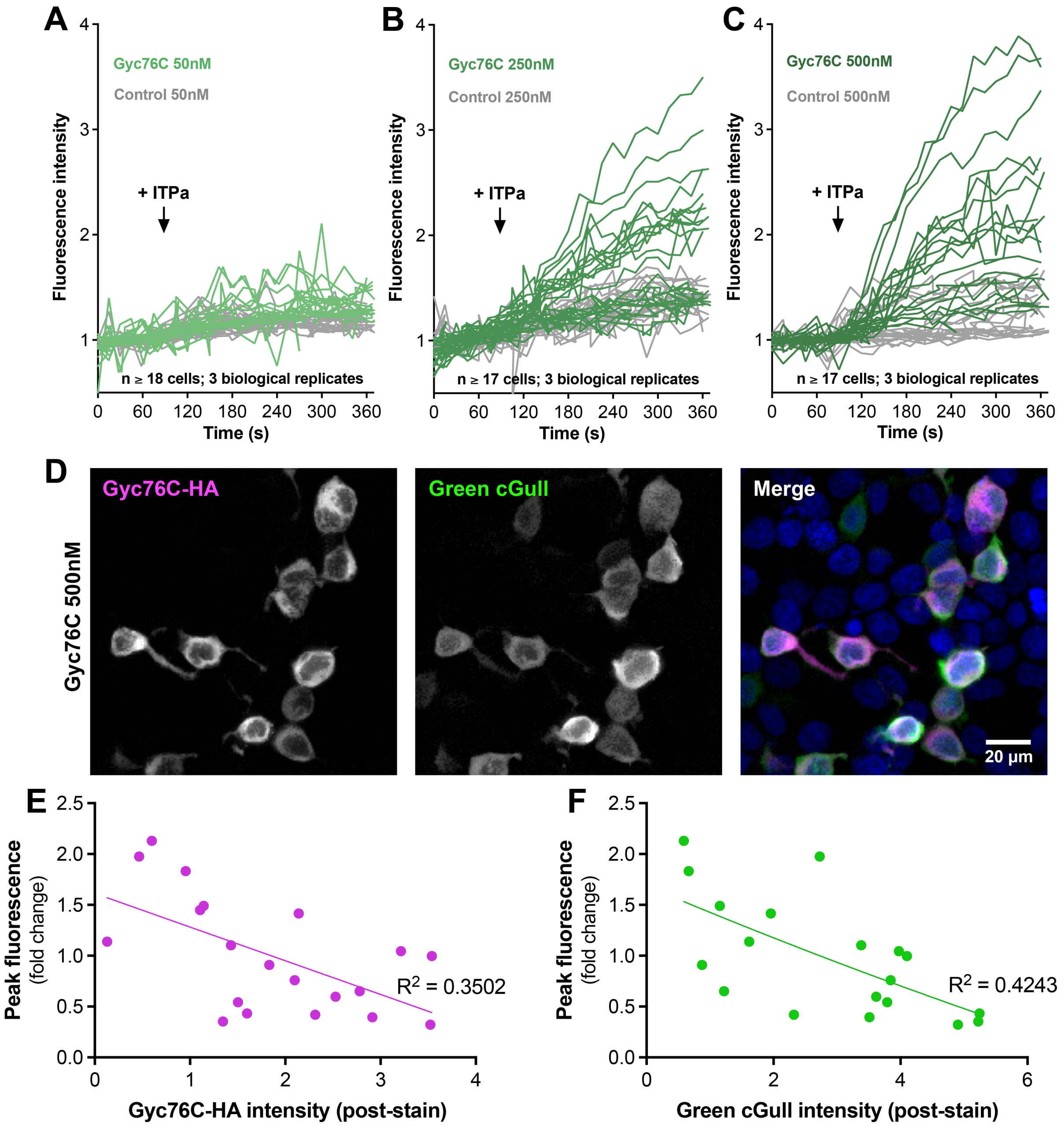


**Figure 7 Supplement 1: ITPa activates Gyc76C in HEK293T cells.** Application of **(A)** 50nm, **(B)** 250nm and **(C)** 500nM *Drosophila* ITPa to HEK293T cells transiently expressing Green cGull (cGMP sensor) and Gyc76C (green) results in an increase in fluorescence compared to control cells (grey) which do not express Gyc76C. The graph represents the fluorescent intensities of individual cells over time. **(D)** Post-hoc immunohistochemistry of tested HEK293T cells (500nM ITPa) reveals that not all cGull expressing cells express Gyc76C-HA, explaining the lack of response to ITPa in some cells. **(E and F)** Graphs depicting stained Gyc76C-HA (E) and Green cGull (F) fluorescence intensities plotted against peak live Green cGull fluorescence increases (500nM condition) indicate weak negative correlations between exogenous protein expression level and change in biosensor fluorescence.

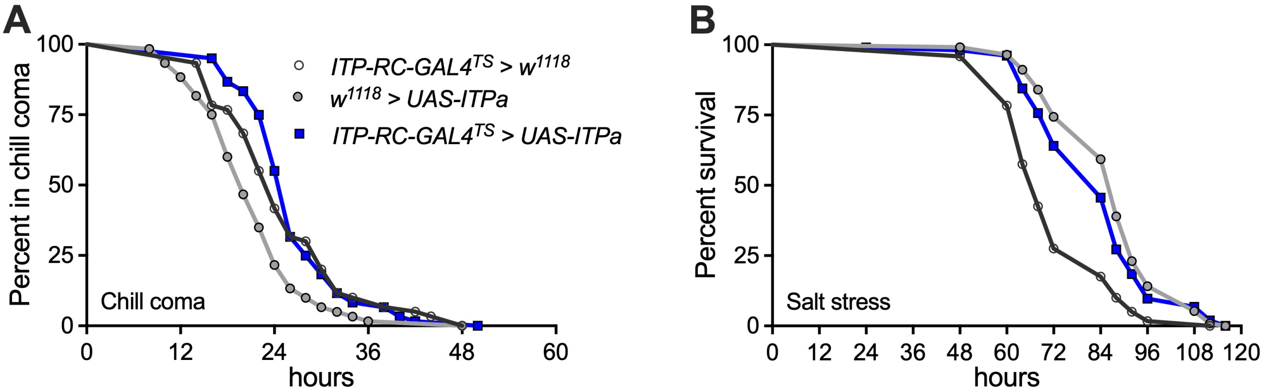


**Figure 10 Supplement 1: ITPa overexpression in adult female *Drosophila* has no effect on cold and ionic stress.** Overexpression of ITPa using *ITP-RC-GAL4$^{TS}$* has no impact on **(A)** recovery from chill coma and **(B)** tolerance to salt stress, as assessed by Log-rank (Mantel-Cox) test.

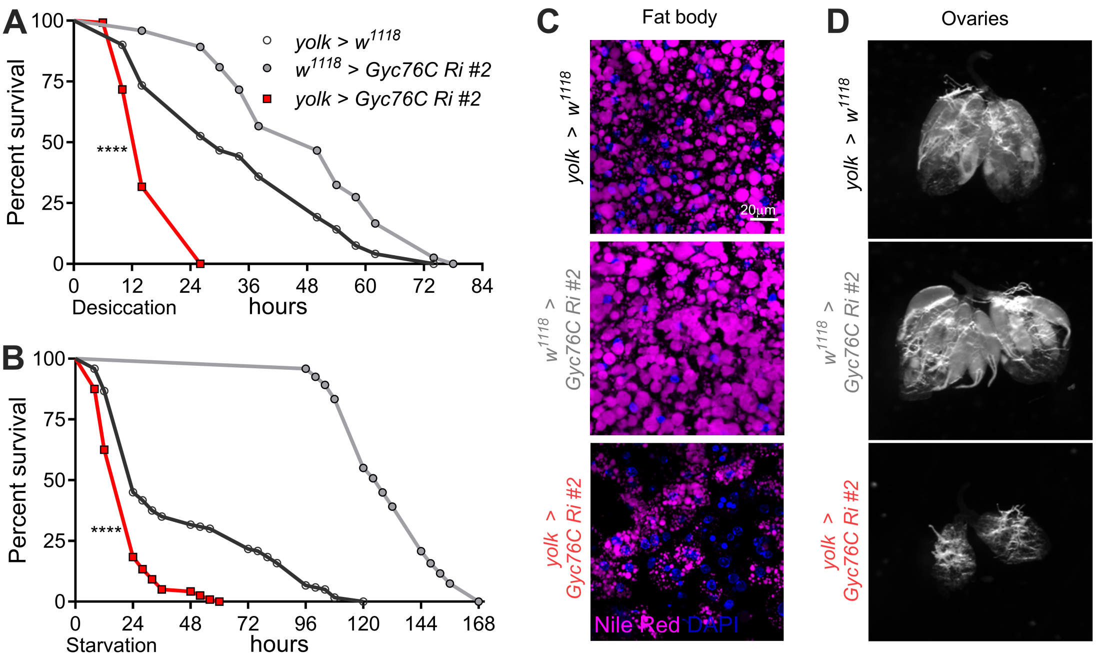


**Figure 12 Supplement 1: *Gyc76C* knockdown with an independent RNAi in females using *yolk-GAL4* impacts stress tolerance, energy stores and reproductive physiology.** Flies with fat body specific *Gyc76C* knockdown using *UAS-Gyc76C RNAi #2* (*#57315*) are **(A)** extremely susceptible to desiccation as well as **(B)** starvation. **(C)** Furthermore, flies with *Gyc76C* knockdown in the fat body have **(C)** reduced lipid levels (Nile Red), and **(D)** smaller ovaries. Abbreviation: *Gyc76C Ri #2, Gyc76C RNAi #2*. For **A** and **B**, **** $p < 0.0001$, as assessed by Log-rank (Mantel-Cox) test.

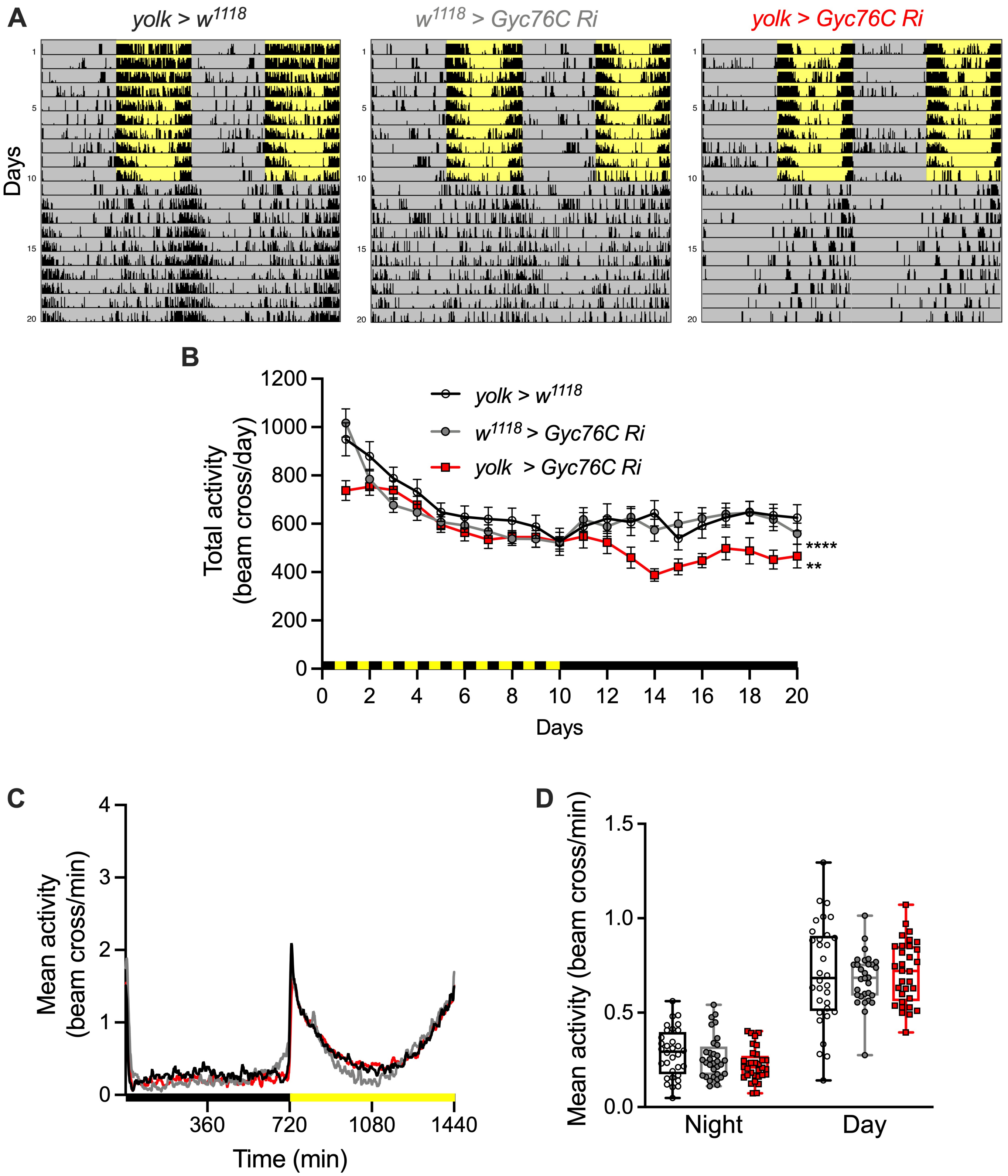


**Figure 12 Supplement 2: *Gyc76C* knockdown with *UAS-Gyc76C Ri* in the female fat body impacts general locomotor activity. (A)** Representative actograms (double-plotted) showing locomotor activity over 20 days. Yellow shading indicates light and grey shading indicates dark conditions. **(B)** Daily total activity of flies over 20 days. *Gyc76C* knockdown flies have reduced locomotor activity on the first day as well as under constant darkness. ** $p < 0.01$ and **** $p < 0.0001$ as assessed by repeated measures one-way ANOVA followed by Tukey's multiple comparisons test. **(C)** Average activity profiles over days 2 to 6. **(D)** Average night- and day-time activities between days 2 to 6 were not significantly different. Abbreviation: *Gyc76C Ri, Gyc76C RNAi.*

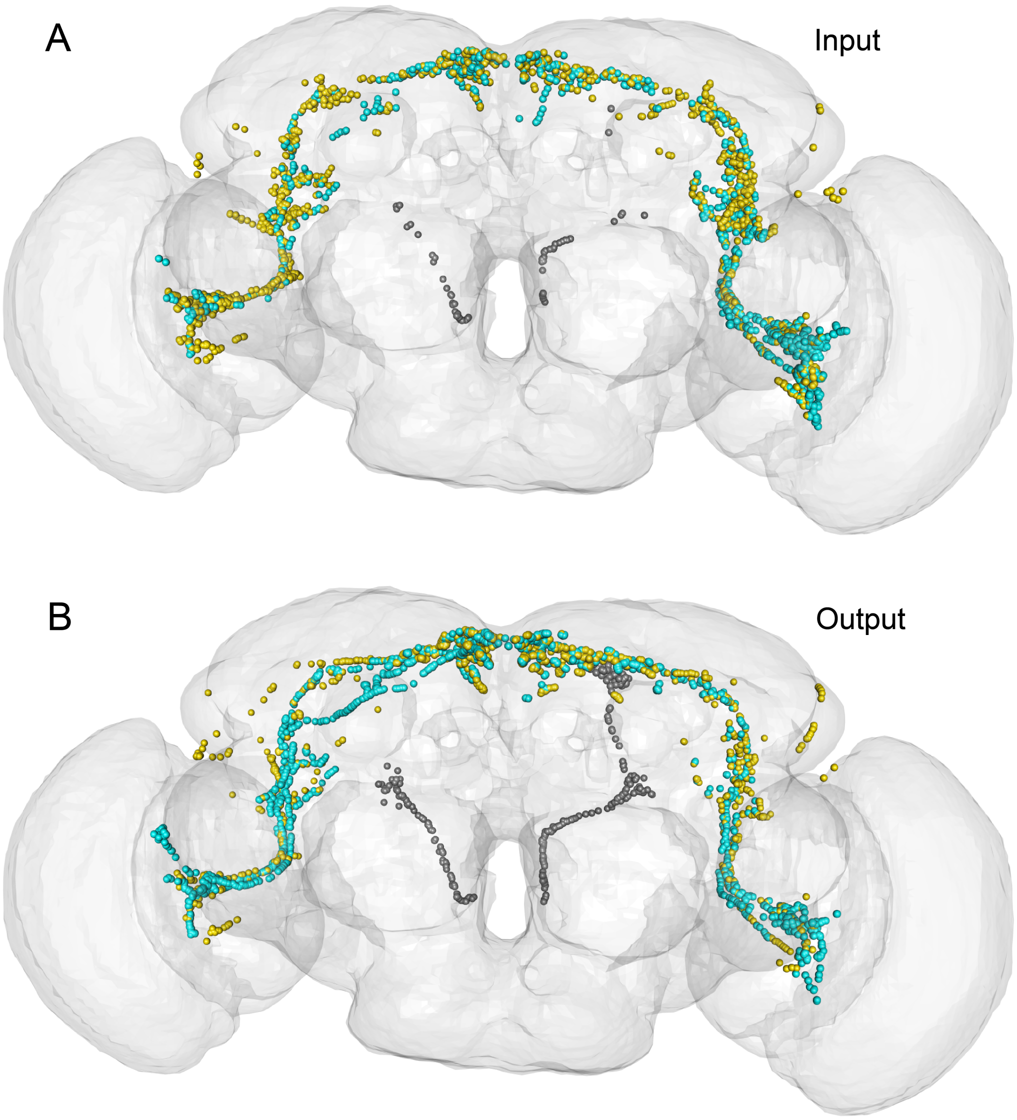


**Figure 13 Supplement 1: Input and output synapses of ITP neurons. (A)** Input and **(B)** output synapses of L-NSC$^{ITP}$ (grey), LN$_d^{ITP}$ (yellow) and 5$^{th}$-LN$_v$ (cyan). Higher magnification of images shown in **Fig. 13B**. All synapses, including those not contributing towards significant connections (less than 5 synapses per connection), are shown here.

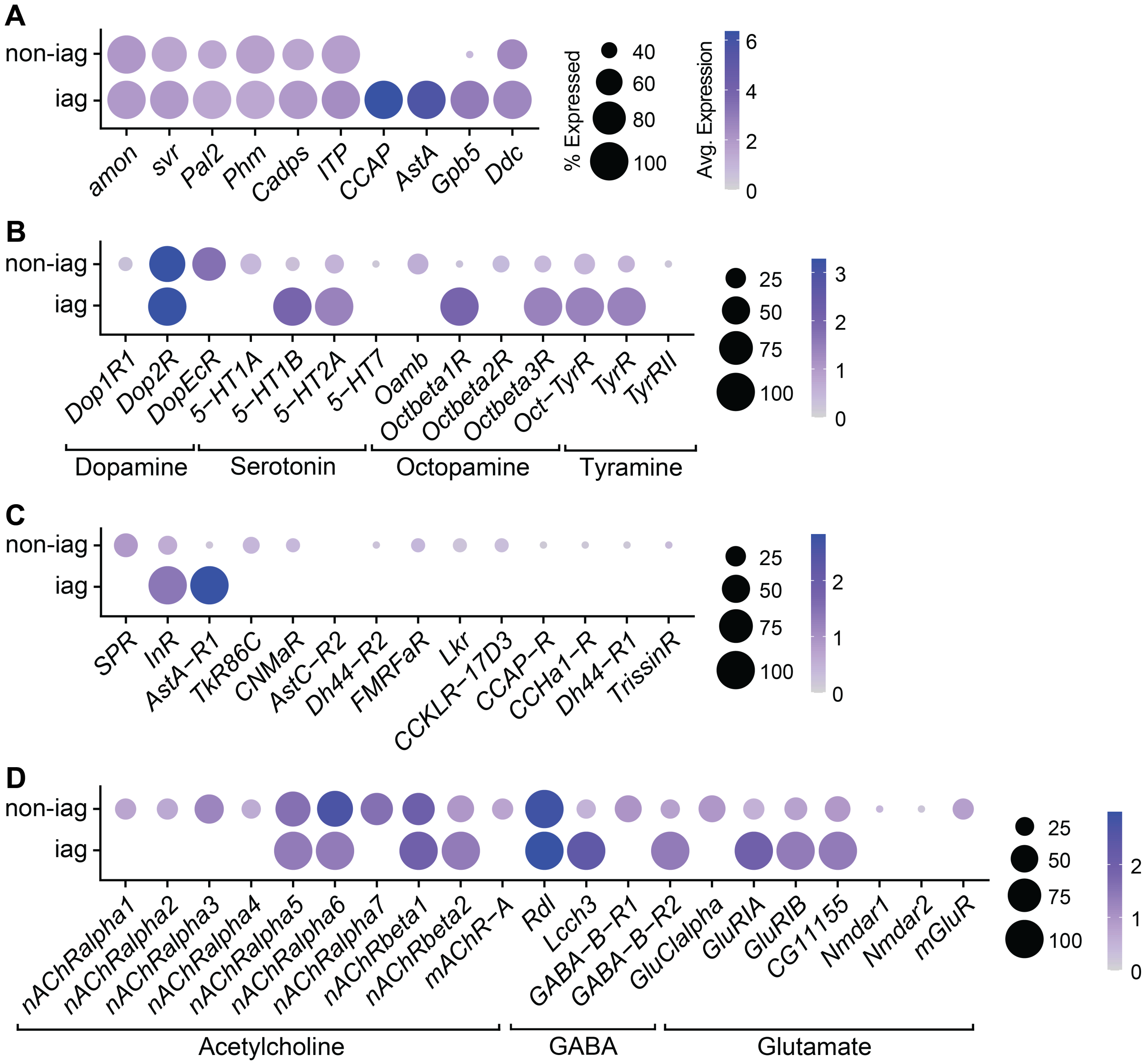


**Figure 13 Supplement 2: Single-cell transcriptomes of *ITP* neurons in the ventral nerve cord.** Identification of single-cell transcriptomes representing ITPa-expressing efferent neurons in the abdominal ganglion (iag) and other *ITP* neurons (non-iag) in the ventral nerve cord dataset (Allen *et al.*, 2020). **(A)** Both sets of neurons express genes required for neuropeptide processing and release (*amon*, *svr*, *Pal2*, *Phm* and *Cadps*) and were identified based on the neuropeptides (*ITP*, *CCAP*, *AstA*) they express. Dot plots showing expression of **(B)** monoamine, **(C)** neuropeptide and **(D)** neurotransmitter receptors in iag and non-iag neurons.
